# Immunity to SARS-CoV-2 up to 15 months after infection

**DOI:** 10.1101/2021.10.08.463699

**Authors:** Harold Marcotte, Antonio Piralla, Fanglei Zuo, Likun Du, Irene Cassaniti, Hui Wan, Makiko Kumagai-Braesh, Juni Andréll, Elena Percivalle, Josè Camilla Sammartino, Yating Wang, Stelios Vlachiotis, Janine Attevall, Federica Bergami, Alessandro Ferrari, Marta Colaneri, Marco Vecchia, Margherita Sambo, Valentina Zuccaro, Erika Asperges, Raffaele Bruno, Tiberio Oggionni, Federica Meloni, Hassan Abolhassanni, Federico Bertoglio, Maren Schubert, Luigi Calzolai, Luca Varani, Michael Hust, Yintong Xue, Lennart Hammarström, Fausto Baldanti, Qiang Pan-Hammarström

**Author notes:** These authors contributed equally to this work (Harold Marcotte, Antonio Piralla, Fanglei Zuo, Likun Du). Co-senior authors (Yintong Xue, Lennart Hammarström, Fausto Baldanti, Qiang Pan-Hammarström).

## Abstract

**Background:** Information concerning the longevity of immunity to severe acute respiratory syndrome coronavirus 2 (SARS-CoV-2) following natural infection may have considerable implications for durability of immunity induced by vaccines. Here, we monitored the SARS-CoV-2 specific immune response in convalescent coronavirus disease-2019 (COVID-19) patients up to 15 months after symptoms onset.

**Methods:** The levels of anti-spike and anti-receptor binding domain antibodies and neutralizing activities were tested in a total of 188 samples from 136 convalescent patients who experience mild to critical COVID-19. Specific memory B and T cell responses were measured in 76 peripheral blood mononuclear cell samples collected from 54 patients. Twenty-three vaccinated individuals were included for comparison.

**Findings:** Following a peak at day 15-28 post-infection, the IgG antibody response and plasma neutralizing titers gradually decreased over time but stabilized after 6 months. Plasma neutralizing activity against G614 was still detected in 87% of the patients at 6-15 months. Compared to G614, the median neutralizing titers against Beta, Gamma and Delta variants in plasma collected at early (15-103 days) and late (9-15 month) convalescence were 16- and 8-fold lower, respectively. SARS-CoV-2-specific memory B and T cells reached a peak at 3-6 months and persisted in the majority of patients up to 15 months although a significant decrease in specific T cells was observed between 6 and 15 months.

**Conclusion:** The data suggest that antiviral specific immunity especially memory B cells in COVID-19 convalescent patients is long-lasting, but some variants of concern, including the fast-spreading Delta variant, may at least partially escape the neutralizing activity of plasma antibodies.

**Funding:** EU-ATAC consortium, the Italian Ministry of Health, the Swedish Research Council, SciLifeLab, and KAW.

## INTRODUCTION

Coronavirus disease 2019 (COVID-19), caused by the novel severe acute respiratory syndrome coronavirus 2 (SARS-CoV-2), rapidly resulted in a pandemic constituting a global health emergency. The COVID-19 pathological process exhibits a wide spectrum of clinical manifestations, ranging from asymptomatic to mild, moderate, severe and critical disease. The genome of SARS-CoV-2 encodes four major structural proteins that occur in all coronavirus species: spike protein (S), nucleoprotein (N), membrane protein (M), and envelope protein (E)^1^. The S protein binds to the host receptor (Angiotensin Converting Enzyme 2 [ACE2]) through the receptor-binding domain (RBD) in the S1 subunit, followed by the S2 subunit-mediated cell membrane fusion^2^.

The adaptive immune response is likely to be critical for the development of protective immunity to SARS-CoV-2 including viral clearance and the persistence of antiviral immunity^25^. Generation of neutralizing antibodies that specifically target the receptor-binding domain (RBD) of the S protein is considered to be essential in controlling SARS-CoV-2 infection^3,4^. A robust adaptive immune response with presence of RBD and S-specific neutralizing antibodies, memory B cells and T cell response have been found in patients who have recovered from infection^5–7^. Although circulating antibodies derived from plasma cells wane over time, long-lived immunological memory can persist in expanded clones of memory B cells^7^.

Since December 2020, variants of concern (VOCs), which were initially named based on the country where they were first identified, have been characterized, including Alpha (B.1.1.7, UK), Beta (B.1.351, South Africa), Gamma (P.1, Brazil) and Delta (B.1.617.2, India). Mutations in the S1 subunit may lead to changes in the structure of the S protein and RBD^8^ and result in higher binding of the virus to the receptor^9^, increased risk of transmission and severity of illness^10^, as well as a reduction in neutralization susceptibility by antibodies^11–13^. The Delta variant is associated with an estimated 60% higher risk of household transmission than the Alpha variant and is becoming dominant worldwide^14^.

We have previously reported the longevity of the SARS-CoV-2 adaptive immune response (up to 6-8 months) in cohorts of Swedish and Italian patients infected with the SARS-CoV-2 G614 strain^6^. The antibody and neutralizing titers were sustained at a relatively high level for at least 6 months after the onset of symptoms while specific memory B and T cells were maintained for at least 6-8 months. In this study, the adaptive immune response in convalescent patients from the same cohorts was followed up to 15 months. In addition, the specific antibody levels and neutralizing antibody titers were tested against VOCs.

## Results

### Longevity of anti-SARS-CoV-2 antibody response

For the entire 15 months follow-up, a total of 188 serum or plasma samples were collected from 136 COVID-19 patients (98 from Italy and 38 from Sweden) experiencing mild symptoms to critical disease (Figure 1, Table S1 and Table S2). Plasma from 108 historical negative controls collected before the SARS-CoV-2 pandemic were also analyzed. Plasma anti-RBD and anti-S antibody titers were measured by an in-house enzyme-linked immunosorbent assay (ELISA)^6^.

**Figure 1.**
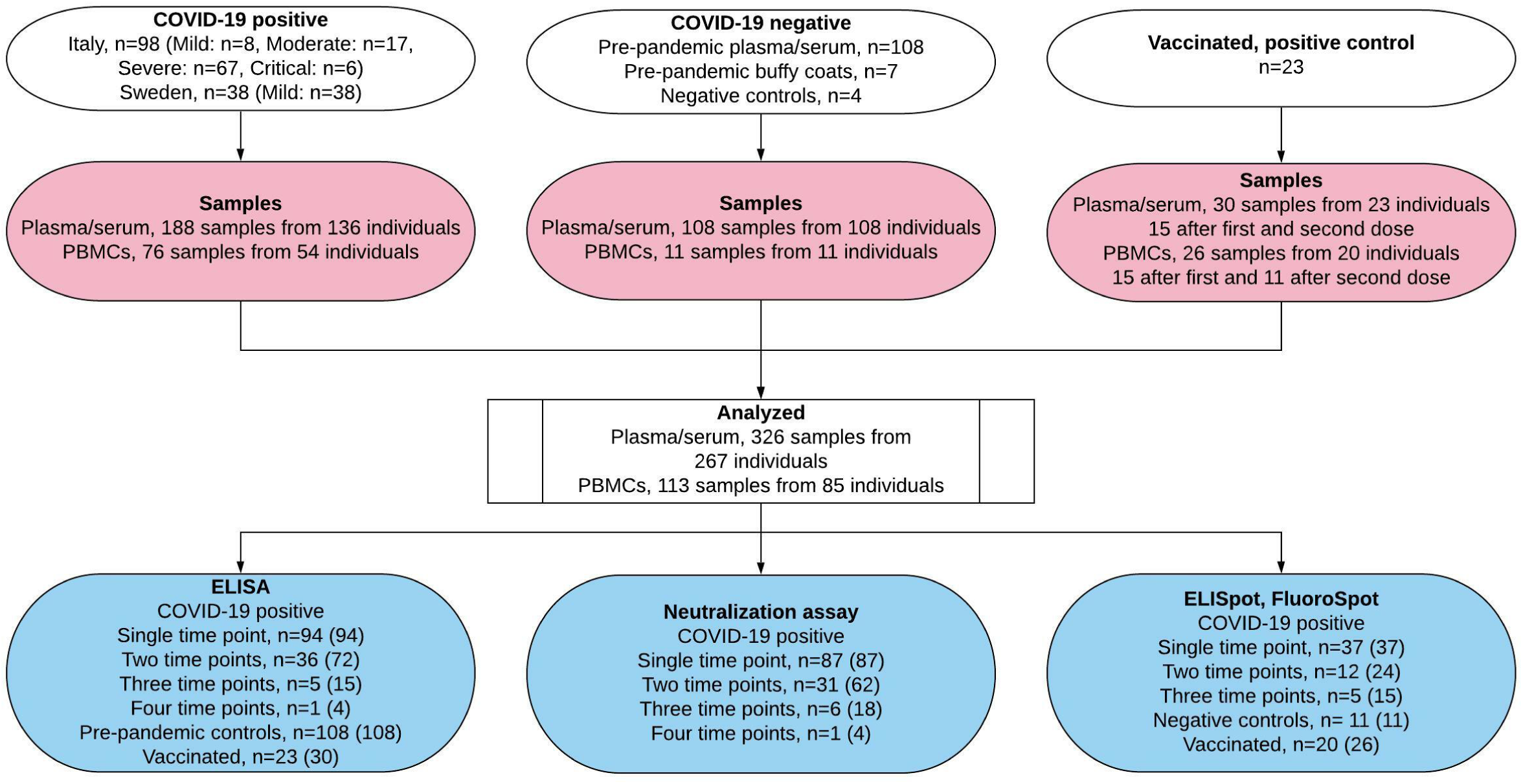
Flowchart illustrating the study design and analysis. For ELISA, Neutralization and ELISpot/Fluorospot, number of individuals (n =) and the number of analyzed samples (in parenthesis) are indicated. See also Table S1-S3.

At the peak of the antibody response, 15-28 days after symptoms onset, anti-RBD IgM and IgA were increased in 77% (40/52) and 85% (44/52) of convalescent patients, respectively, but rapidly decreased between 1 and 3 months and were detected in less than 4.5% (2/44) and 11% (5/44) of patients tested between 6 and 15 months when assessing all COVID-19 subjects by cross-sectional analysis (Figures 2A, 2B, 2D, 2E). Similarly, IgM and IgA anti-S proteins were detected in 88% (46/52) and 90% (47/52) of convalescent patients at 15-28 days, respectively, but less than 23% (10/44) of patients for both immunoglobulins from 6 to 15 months (Figures 2G, 2H, 2J, 2K). Using a one-phase exponential decay model, we estimated the half-lives (t_1/2_) of the RBD- and S-specific IgM antibodies to be 55 and 65 days, respectively, (Figures 2A, 2G), and that of RBD- and S-specific IgA antibodies to be 56 days and 55 days, respectively (Figures 2B, 2H).

**Figure 2.**
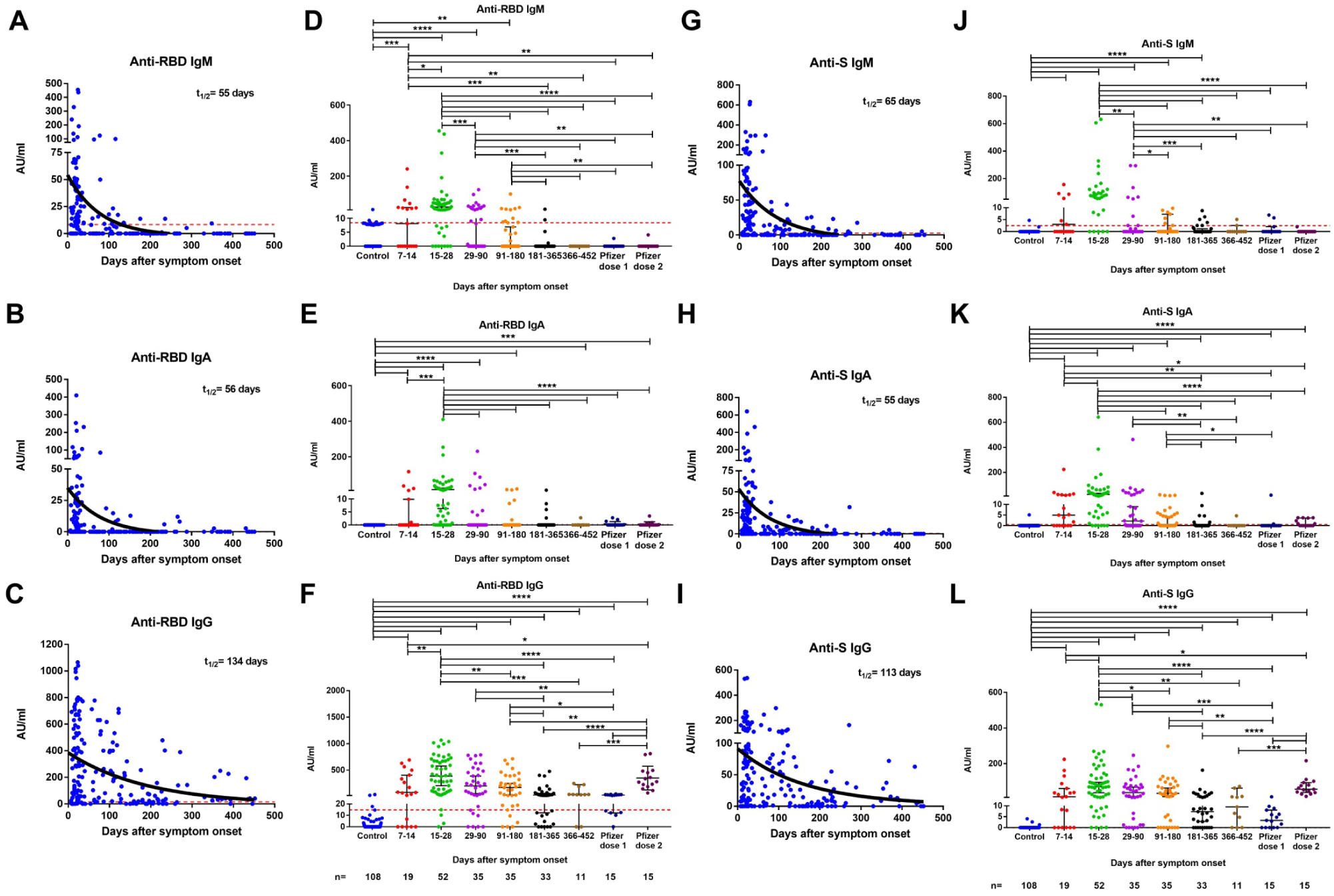
Cross-sectional analysis of plasma anti-SARS-CoV-2 antibody titers patients over time. Levels of anti-RBD (A-F) and anti-S (G-L) IgM, IgA and IgG antibodies in plasma of COVID-19 patients, historical controls, and vaccinated individuals. Antibodies were measured in 185 samples from 136 COVID-19 patients, 108 historical controls (before the SARS-CoV-2 pandemic) and 23 vaccinated individuals. The RBD (A-C) and S (G-I) specific IgM, IgG, IgA antibody decay curves (in black) and half-lives (t^1/2^) were estimated by a one-phase exponential decay model. Samples from patients were further divided in 7 study periods: 7-14 days (n = 19), 15-28 days (n = 52), 29-90 days (n = 35), 91-180 days (n = 35), 181-365 days (n = 33) and 366-452 (n = 11) after symptom onset (D-F and J-L) for comparison. Vaccinated individuals were sampled 14-35 days after the first dose and 14-36 days after the second dose. Symbols represent individual subjects; horizontal black lines indicate the median and 95% confidence interval. The dashed red line indicates the cutoff value for elevated anti-S and anti-RBD antibody titers (2.5 and 8.4 AU/mL for IgM, 0.5 and 0.08 AU/mL for IgA, and 0.03 and 14.81 AU/mL for IgG, respectively). Mann-Whitney U test. *p ≤ 0.05, **p ≤ 0.01, ***p ≤ 0.001, and ****p < 0.0001. See also Figures S1 and S2.

Plasma IgG antibodies binding to SARS-CoV-2 RBD and S protein increased in 94% (49/52) of COVID-19 convalescent participants tested 15-28 days after symptoms onset. The median RBD and S IgG antibody titers gradually decreased by less than 4-fold from the peak of the antibody response until 6 months (15-28 vs 181-365 days, p < 0.0001 for both antibodies), but thereafter, remained at a relatively steady level up to 15 months (181-365 vs 366-452 days, p = 0.3866 and p = 0.7105 for anti-RBD and anti-S antibody titers, respectively; Figures 2C, 2F, 2I, 2L). Both anti-RBD and anti-S IgG antibodies were still detected in 68% (30/44) and 73% (32/44) of patient samples tested 6 to 15 months after symptoms onset (Figures 2F, 2L). The half-lives of anti-RBD and anti-S IgG antibody response estimated by a one-phase decay model were 134 and 113 days, respectively (Figures 2C, 2I) and were shorter in patients with mild/moderate (t_1/2_ = 52 days and t_1/2_ = 40 days) than severe/critical (t_1/2_ = 372 days and t_1/2_ = 239 days) disease (Figures S1A and S1B).

As a complementary approach, we analyzed the antibody titers from 42 patients who donated blood at two or more time points and estimated the half-lives of the RBD- and S-specific antibody response in IgM (t_1/2_ = 71 days and 73 days), IgA (t_1/2_ = 32 days and 28 days) and IgG (t_1/2_ = 128 days and 90 days) (longitudinal analysis; Figure 3). No significant difference in the anti-S and anti-RBD IgG titers were observed between the paired samples (n = 11) containing a sampling time point between 181 and 300 days (6-10 months) and a second one between 301 and 452 days (10-15 months), confirming that specific IgG antibody titers reached a plateau phase after 6 months (p = 0.8984 and p= 0.3125, respectively; Figures 3C and 3F).

**Figure 3.**
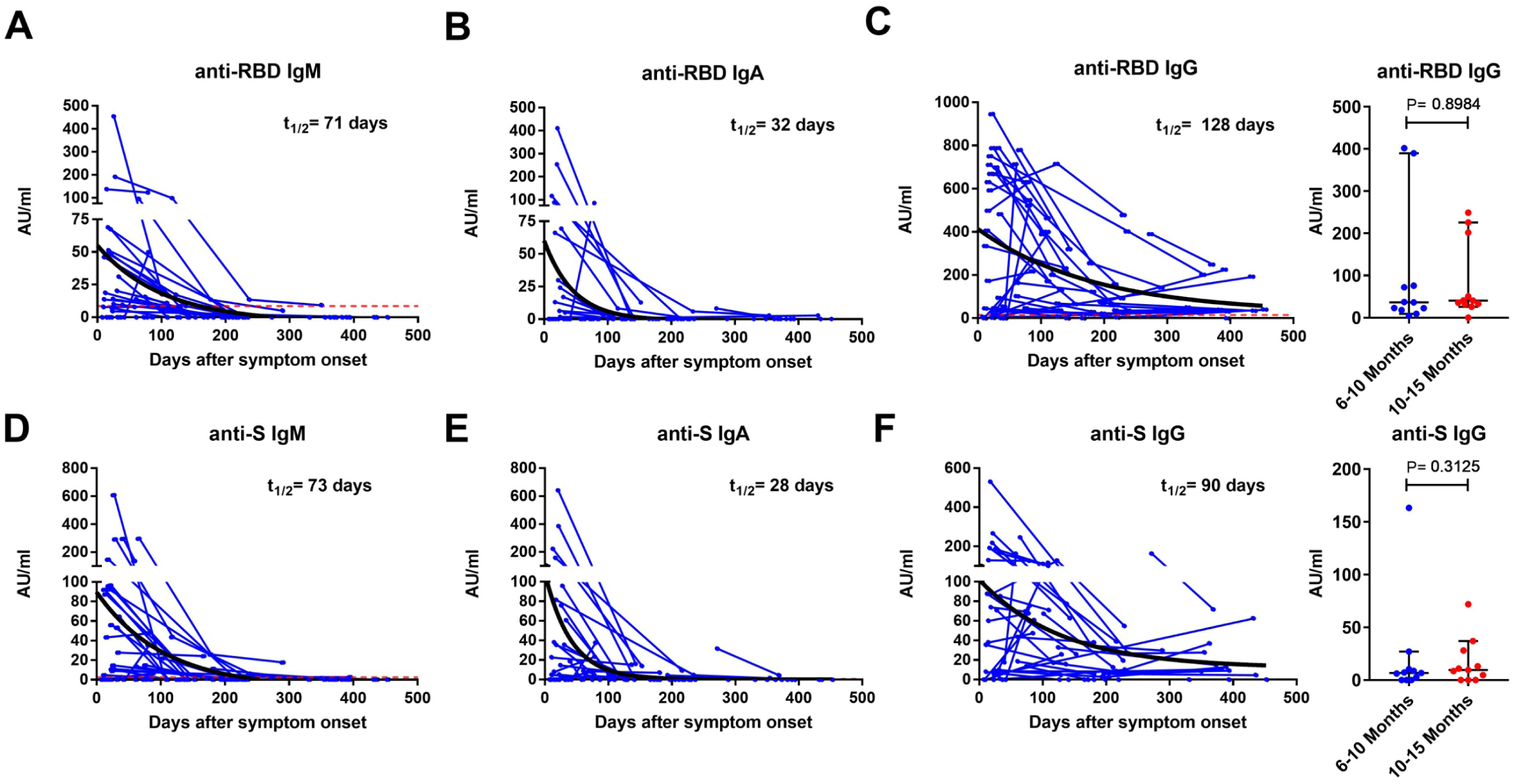
Longitudinal analysis of plasma antibody titers. Anti-RBD (A-C) and anti-S (D-F) IgM, IgA and IgG antibodies in paired samples from 42 COVID-19 patients over 15 months. For anti-RBD (C) and anti-S (F) IgG antibodies, further comparisons were made between paired samples (n = 11) at two time points ranging from 6 to 15 months (TP1: 181-300 and TP2: 301-452 days after symptoms onset; right panel). Symbols represent individual subjects; horizontal black lines indicate the median and 95% confidence interval. The antibody decay curves (in black) and half-lives (t^1/2^) were estimated by a one-phase exponential decay model. Wilcoxon signed-rank test. See also Figure S1.

We further compared the antibody response induced from natural infection to that induced by one or two doses of the Comirnaty (Pfizer-BioNTech) vaccine (Table S3). The vaccine induced no or low level of RBD- and S-specific IgM (Figures 2D, 2J) and IgA (Figures 2E, 2K) antibody titers compared to that detected during natural infections. The plasma RBD and S specific IgG antibody titers measured 14-35 days after one dose of vaccine were similar to those measured in patients more than six months after infection when the antibody titers have decreased (vs 181-365 days, p = 0.6959 and p = 0.1169, respectively; vs 366-452 days, p = 0.0685 and p = 0.1121, respectively; Figures 2F and 2L). The RBD IgG and S antibody titers measured 14-36 days after the second dose of vaccine corresponded to those observed at the peak of antibody response in convalescent patients (15-28 days) (p = 0.7489 and p = 0.8435, respectively; Figures 2F and 2L).

The neutralizing activity against the G614 variant was measured by a microneutralization assay and was expressed as the neutralizing titers (≥ 1:10) which inhibit 90% of the virus infectivity (NT_90_). Similar to the dynamic of the anti-RBD and anti-S antibodies, the median NT_90_ reached a peak of 1:160 between 15-28 days with 98% (51/52) of convalescent patients exhibiting plasma neutralizing activity (Figures 4A and 4B). The plasma NT_90_ gradually decreased by about 2-fold up to 6 months (15-28 vs 91-180 days, p = 0.0493) but was still observed in more than 87% (35/40) of the patients sampled at 181-365 (27/31) and 366-452 days (8/9) (Figure 4B). Furthermore, no significant difference was observed in the median NT_90_ measured at 181-365 (1:40) and 366-452 days (1:80) (p = 0.6674; Figure 4B) or between paired samples at two time points between 6 and 15 months (n = 9, TP1: 181-300 days vs TP2: 301-452 days, p = 0.5156; Figure 4C). A one-phase decay model showed a rapid initial decay using both cross-sectional (t_1/2_ = 70 days, Figure 4A) or longitudinal analysis (t_1/2_ = 44 days, Figure 4C) which slow down to a plateau phase extending from around 4 up to 15 months. A decrease of NT_90_ followed by a plateau phase was observed in both patients with mild/moderate and severe/critical diseases using cross-sectional analysis (Figure S1C) while no decline was observed for the severe/critical patients group using longitudinal analysis (Figure S1F) suggesting that neutralization activity may persist longer in this group. The NT_90_ directly correlated (p < 0.0001) with the levels of RBD-specific IgM (r = 0.45), IgA (r = 0.37) and IgG (r = 0.49) as well as with the level of S-specific IgM (r = 0.44), IgA (r = 0.44) and IgG (r = 0.55) antibodies (Figure S2).

**Figure 4.**
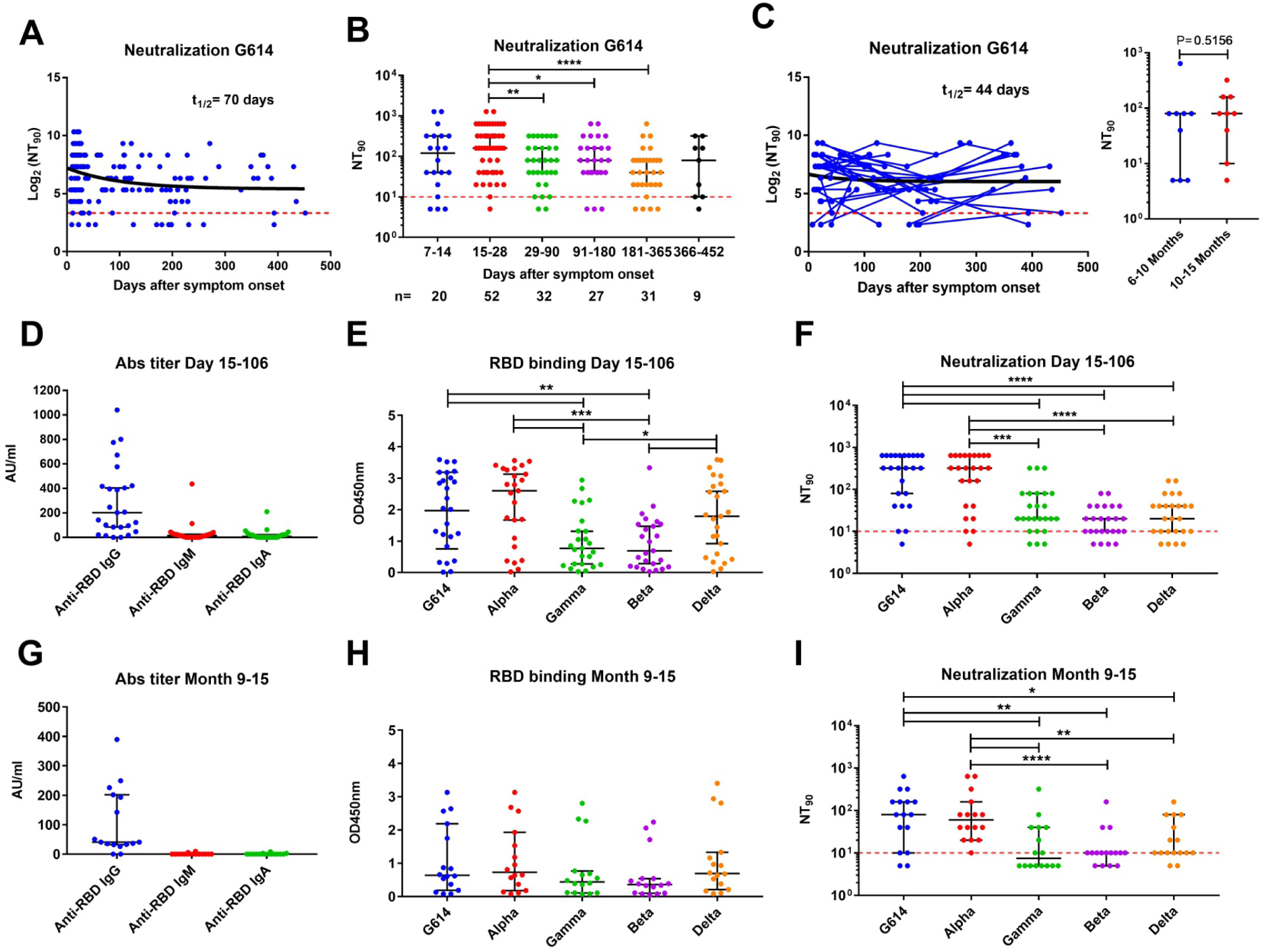
Cross-sectional and longitudinal analysis of plasma neutralization activity against SARS-CoV-2 and variants of concern. (A) Dynamics of plasma neutralizing activity against G614 variant in COVID-19 patient samples over time. (B) Samples from patients were taken at 7 study periods: 7-14 days (n = 20), 15-28 days (n = 52), 29-90 days (n = 32), 91-180 days (n = 27), 181-365 days (n = 31) and 366-452 (n = 9) after symptom onset. (C) For longitudinal analysis, samples were taken at two (n = 31) or more (n = 7) time points, and further comparisons were made between paired samples (n = 9) at two time points ranging from 6 to 15 months (TP1: 181-300 and TP2: 301-452 days after symptoms onset; right panel). The NT_90_ decay curves (in black) and corresponding half-lives (t_1/2_) were estimated by a one-phase decay model (A, C). To test cross-neutralization, the level of anti-RBD IgM, IgA, and IgG titers (D, G), binding activity of IgG antibody to RBD from SARS-CoV-2 variants (E, H), and plasma neutralizing activity against variants (F, I) were tested in plasma collected from COVID-19 patients at 15-106 days (median day of 24) and 9-15 months (241-452 days, median day of 370). The dashed red line indicates the titer cutoff value (≥ 1:10). Symbols represent individual subjects; horizontal black lines indicate the median and 95% confidence interval. Mann-Whitney U test. *p≤ 0.05, **p ≤ 0.01, ***p ≤ 0.001 and ****p < 0.0001. See also Figures S1 and S2.

In order to evaluate if the convalescent patients could be protected from the circulating VOCs, we determined the cross-binding and cross-neutralizing activity against Alpha, Gamma, Beta and Delta variants using plasma collected at the early (15-106 days, median day of 24) and late (259-452 days, median day of 370) phase of convalescence from February to May 2021. As described earlier, the IgG anti-RBD titers against wild-type RBD were higher in early (Figure 4D) compared to late convalescence (Figure 4G). Furthermore, a reduction in the level of IgG antibodies binding to Gamma and Beta RBD was observed using both plasma samples collected at 15-106 days and 259-452 days (9-15 months) although it was only significant in early convalescence (15-106 days, p = 0.0039 and p = 0.0045 for Gamma and Beta, respectively; Figures 4E and 4H). The majority of samples (96%, 24/25) collected at earlier time points (15-106 days) showed neutralizing activity against G614 and Alpha, and 84-88% of them against the other VOCs. The median NT_90_ was similar (1:320) against G614 and Alpha but 16-fold lower (1:20, p < 0.0002) for Gamma, Beta, and Delta variants (Figure 4F). Between 9 to15 months after infection, the majority of patient samples still showed neutralizing activity against G614 (14/16, 88%), Alpha (16/16, 100%), Beta (12/16, 75%) and Delta (14/16, 88%) variants, but a lower proportion of patient samples had neutralizing activity against Gamma (8/16, 50%) (Fisher exact test, p = 0.002 compared to Alpha; Figure 4I). The NT_90_ were 6- to 8-fold lower against Gamma (< 1:10, p = 0.0039 and p = 0.0010), Beta (1:10, p = 0.0025 and p < 0.001) and Delta (1:10, p = 0.0237 and p = 0.0050) compared to G614 (1:80) and Alpha (1:60), respectively (Figure 4I).

These data suggest that the plasma anti-SARS-CoV-2 antibody response and neutralizing activity decrease up to around 6 months but neutralizing activity is maintained in the majority of patients up to 15 months. Furthermore, plasma neutralizing activity was lower against Beta, Gamma and Delta variants, particularly 9-15 months after infection.

### SARS-CoV-2-specific memory B cells

We analyzed 76 peripheral blood mononuclear cell (PBMC) samples collected from 54 patients (mild = 34, moderate = 4, severe = 15, critical = 1) for the presence of SARS-CoV-2 specific B and T cells. Using the highest value observed from all of the COVID-19 negative controls as a cutoff, RBD-specific IgG-producing B cells were detected by ELISpot assay in 89% (68/76) of the patient samples tested (Figure 5A). We further divided samples into 5 groups based on the sample time points; collected at 14-30 days, 31-90 days, 91-180 days, 181-365 days and 366-452 days, post-onset of symptoms, i.e. ∼2-4 weeks, 1-3 months, 3-6 months, 6-12 months, and 12-15 months, respectively. RBD-specific IgG-producing B cells were detected in 64% (7/11), 78% (7/9), and 90% (18/20) of the samples collected at 2-4 weeks, 1-3 month and 3-6 months, respectively, and in all samples collected at 6-12 (n = 28) and 12-15 (n = 8) months after the onset of symptoms (Figure 5B). After an expansion over the first 3 months, the RBD-specific IgG-producing cells reached a maximum at 3–6 months (91-180 days) post-infection followed by a gradual but non-significant decrease after 6 months using cross-sectional analysis (91-180 vs 181-365 days, p = 0.1518; vs 366-452 days, p = 0.3039; Figure 5B) or by comparing the paired samples at two time points ranging between 6 and 15 months (n = 9, TP1: 180-300 vs TP2: 301-452 days, p = 0.3008; Figure 5C). RBD-specific IgG-producing B cells were still maintained in relatively high numbers in all patients (n = 36) followed up between 6 and 15 months.

**Figure 5.**
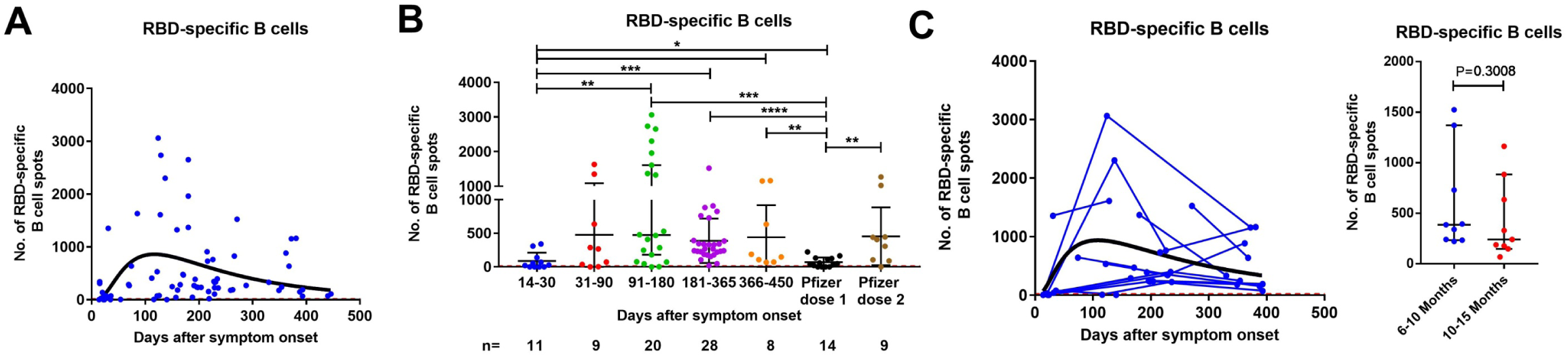
Cross-sectional and longitudinal analysis of SARS-CoV-2-specific memory B cell responses in COVID-19 patients. (A) Dynamics of RBD-specific memory B cells levels in COVID-19 patient samples over time with the corresponding log-normal fitting curve (in black). (B) B cells were measured in control (n = 11), COVID-19 samples at 5 study periods: 14-30 days (n = 11), 31-90 days (n = 9), 91-180 (n = 20), 181-365 (n = 28), 366-452 (n = 8) days after symptom onset, as well as vaccinee samples after first (n = 14) and second (n = 9) dose. For longitudinal analysis (C), samples were taken at two (n = 12) or more (n = 5) time points, and further comparisons were made between paired samples (n = 9) at two time points ranging from 6 to 15 months (TP1: 181-300 and TP2: 301-452 days after symptoms onset) (right panel). The results were expressed as the number of spots per 300,000 seeded cells after subtracting the background spots of the negative control. The horizontal black lines indicate the median value and 95% confidence interval of the group. The cutoff value (dashed red line) was set at the highest number of specific B cell spots for the negative controls (> 12 spots / 300,000 cells). Mann-Whitney U test. **p ≤ 0.01, ***p ≤ 0.001, and ****p ≤ 0.0001. See also Figures S3 and S6.

No statistically significant differences were observed in the number of memory B cells between mild/moderate and severe/critical COVID-19 patients over the period ranging from 6 to 15 months suggesting that the intensity and duration of the B cell response are not dependent on the disease severity (p = 0.5835; Figures S3 and S6).

Non-infected individuals sampled 14-35 days after the first vaccine dose showed a B cell response significantly lower than that observed in convalescent patients between 3 and 15 months (vs 91-180 days, p = 0.0002; vs 181-365 days, p < 0.0001; vs 366-452 days, p = 0.0077 after onset of symptoms; Figure 5B). Subjects sampled 14-36 days after the second vaccine dose showed a median number of circulating RBD-specific memory B cells similar to that observed in convalescent patients sampled during the same period (vs 91-180 days, p = 0.3222; vs 181-365 days, p = 0.6337; vs 366-452 days, p = 0.8884; Figure 5B).

### SARS-CoV-2-specific memory T cells

The S1 and S N M O peptide pool-specific T cells expressing interleukin-2 (IL-2) and/or interferon-gamma (IFN-γ) were measured by FluoroSpot assay. No or a negligible number of IL-2, IFN-γ, or IL-2/IFN-γ -producing T cells against the two peptide pools were detected in the negative controls. Overall, a T cell response against at least one of the SARS-CoV-2 peptide pools (S1, or S N M O protein derived) was detectable at a level above the cutoff in 95% (69/73) of the patient samples tested over the study period (Figures 6A-6C and S4A-S4C). When divided by groups based on the sample time points, specific T cells were detected in the majority of patient samples at 2-4 weeks (10/11, 91%) and 1-3 months (7/9, 78%), and more than 98% (52/53) of patient samples tested between 3 and 15 months.

**Figure 6.**
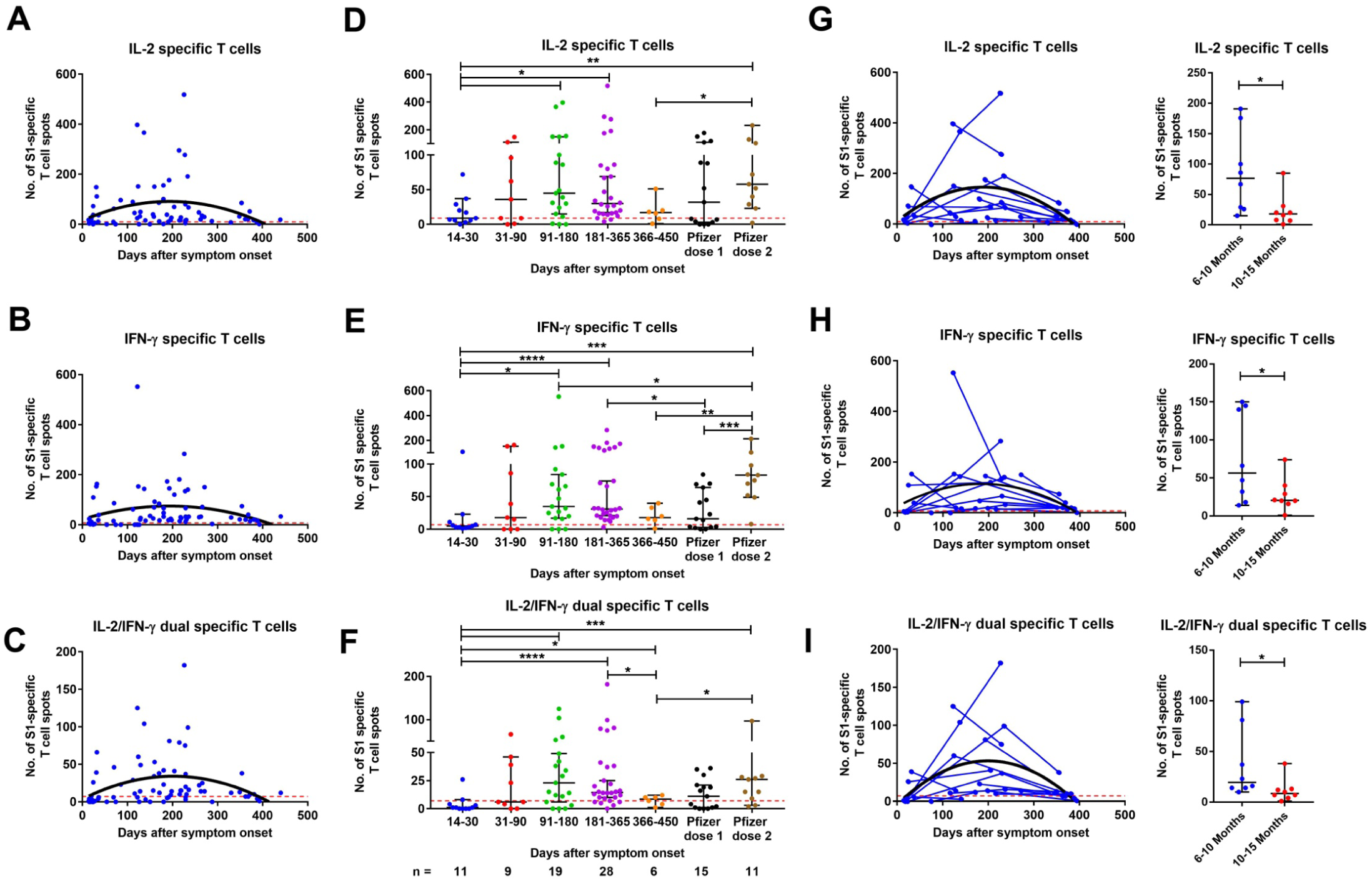
Cross-sectional and longitudinal analysis of S1-specific memory T cell responses in COVID-19 patients. (A-C) Dynamics of S1-specific memory IL-2, IFN-γ, and IL-2/IFN-γ-producing T cells with the corresponding second order polynomial fitting curve (in black). (D-F) T cells were measured in control (n =11), COVID-19 samples at 5 study periods: 14-30 days (n = 11), 31-90 (n = 9), 91-180 days (n = 19), 181-365 (n = 28), and 366-452 (n = 6) days after symptom onset, as well as vaccinee samples after first (n = 15) and second (n = 11) dose. (G-I) For longitudinal analysis, samples were taken at two (n = 10) or more (n = 5) time points, and further comparisons were made between paired samples (n = 8) at two time points ranging from 6 to 15 months (TP1: 181-300 and TP2: 301-452 days after symptoms onset; right panel). The results were expressed as the number of spots per 300,000 seeded cells after subtracting the background spots of the negative control. The horizontal black lines indicate the median value and 95% confidence interval of the group. The cutoff value (dashed red line) was set at the highest number of specific T cell spots for the negative controls (> 7 to 9 spots / 300 000 seeded cells depending on the T cell population). Mann-Whitney U test. **p ≤ 0.01, ***p ≤ 0.001, and ****p ≤ 0.0001. See also Figures S4-S6.

Using cross-sectional analysis, the number of S1-specific IL-2, IFN-γ, and IL-2/IFN-γ-producing T cells reached a peak between 3 and 6 months and was maintained up to 15 months except for IL-2/IFN-γ-producing T cells which significantly decreased at 366-452 days (vs 181-365 days, p = 0.0215; Figures 6D-6F). We further analyzed the results using longitudinal data and observed a significant decrease of S1-specific T cells producing IL-2, IFN-γ, and IL-2/IFN-γ between TP1 (180-300 days) and TP2 (301-452 days) (n = 8; p= 0.0156 for each of the three cell populations; Figures 6G-6I). We similarly detected a peak of S N M O pool-specific IL-2, IFN-γ, and IL-2/IFN-γ-producing T cells between 3 and 6 months and a significant decrease of those T cell populations at 12-15 months using both cross-sectional (366-450 vs 181-365 days; p = 0.0082, p = 0.0004, and p= 0.0026 respectively; Figures S4D-S4F) and paired samples analysis (n = 8, TP1: 181-300 days vs TP2: 301-452 days, p = 0.0078 for the three T cell populations; Figures S4G-S4I).

The numbers of S1 and S N M O peptide pool-specific IL2 and IL-2/IFN-γ -producing T cells were higher in severe/critical COVID-19 than mild/moderate patients between 6 and 15 months (p = 0.0044 and p = 0.0202 for S1 specific T cells, p = 0.0302 and p = 0.0427 for S N M O specific T cells) while no significant difference in S1 and S N M O specific IFN-γ -producing T cells was observed (p = 0.0610 and p = 0.1012, respectively) (Figures S5 and S6).

The S1-specific T cell response measured in samples collected after one vaccine dose was equivalent to that observed in convalescent patients at 12-15 months when the specific T cell number decrease (p = 0.6333, p = 0.8352, p = 0.6893; Figures 6D-6F). Compared to the peak of T cell response in patients (3-6 months), individuals with two doses of vaccine present a similar level of S1-specific IL-2 and IL-2/IFN-γ-producing T cells but a 2-fold higher number of IFN-γ-producing T cells (p = 0.5177, p = 0.9915, and p = 0.0475, respectively; Figures 6D-6F).

Finally, we have investigated the presence of all three arms of immunity for each individual who has been evaluated for SARS-CoV-2-specific adaptive immunity in all assays i.e., 1) IgM, IgG and IgA antibody response and/or neutralization activity, 2) RBD-specific IgG-producing B cells, and 3) S1 peptide or S N M O pool-specific IL-2 and/or IFN-γ-producing T cells (Figure S7). The majority of individuals (72%, 55/76) had three arms of immunity active against SAR-CoV-2, particularly from 2 to 15 months (80%, 48/60), highlighting the increase of memory B and T cells between 0 and 2 months and the long duration of the adaptive immune response.

Thus, SARS-CoV-2-specific memory B and T cells remained present in the majority (>95%) of patients followed up between 6 and 15 months but a reduction of the T cell response was observed at 12-15 months.

## Discussion

The magnitude, duration and quality of immunological memory are crucial for preventing reinfection. In this study, we extended our assessment of the longevity of the SARS-CoV-2-specific antibody, B and T cell immune response in cohorts of convalescent patients in Italy and Sweden that experienced mild to critical symptoms of COVID-19. In accordance with previous studies, although the RBD- and S-specific IgG titers peaked 14-30 days after infection and gradually decreased over time, the IgG antibody response stabilized after 6 months and was still detected in the majority of convalescent plasma donors at 6-15 months^4,5,14,15^. The sustained persistence of RBD-IgG titer over time suggests the generation of long-lived bone marrow plasma cells. Anti-S antibody titers were previously shown to correlate with the frequency of S-specific bone-marrow plasma cells of SARS-CoV-2 convalescent patients 7 to 8 months after infection^17^. Only a low proportion of individuals had anti-S- and anti-RBD IgM or IgA antibodies more than 6 months after infection confirming the faster decrease of the specific IgM and IgA antibody response observed in other studies^5,15,18,19^ although a few studies reported a high prevalence of specific IgA up to one year^7,20^. In accordance with a previous study, the persistence of IgG antibody level was associated with disease severity and patients with milder disease appeared to have more rapid IgG anti-RBD antibody waning^21^. It has been reported that antibodies against SARS-CoV and Middle Eastern respiratory syndrome (MERS)-CoV, could still be detected 1-3 years after infection onset despite lack of re-exposure to this virus^22^. After a rapid decline of antibodies against SARS-CoV in the first two years, specific IgG have been detected in some patients up to 12 years after infection^23^ suggesting that SARS-CoV-2-specific IgG antibodies may be present for a longer period in some individuals.

Functional neutralizing antibodies specific to SARS-CoV-2 (anti-S and anti-RBD) that are produced following infection or vaccination are considered important for viral neutralization and viral clearance^24,25^. In the absence of definitive correlates of protective immunity, the presence of neutralizing antibodies against SARS-CoV-2 provides the best current indication for protection against reinfection^3,4,24^. As previously reported, the neutralizing ability of polyclonal plasma correlated positively with anti-S IgG or anti-RBD IgG^6,19,26^. Plasma neutralizing activity reached a plateau after 4-6 months and was maintained in the majority of patients up to 15 months which was consistent with a previous study showing no significant difference in anti-RBD IgG antibody level and plasma neutralizing activities against Wuhan strain between 6 and 12 months^7^. In addition, the two-phase pattern with an initial rapid decay of neutralizing antibodies which slowed down to a flat slope^27^ might explain why early studies with shorter follow-up (2-4 months) reported a fast decline of the antibody response^28^. The long-lasting neutralizing activity might be caused by the accumulation of somatic mutations in IgG antibody genes and the production of antibodies with increased neutralizing potency^7,27^. Although no significant difference in neutralization activity was observed beyond 6 months, it might be important to predict how the long-term neutralizing antibody decay will affect the clinical outcomes following both natural infection and vaccination. However, a recent study in fully vaccinated health care workers showed that the occurrence of breakthrough infection was more associated with the peak of antibody and neutralization titers induced by the vaccine rather than on the subsequent antibody decay suggesting that the degree of protection might depend more on the initial immune response and the generation of memory B and T cells^24^.

Although there is evidence that SARS-CoV-2 seropositivity is associated with protection against the same strain^29,30^, reinfection has been observed in some patients, which may represent non-durable protective immunity or infection with different variants^31–33^. Notable mutations identified in the S1 subunit of the Alpha (Del69-70, N501Y, P681H), Beta (K417N, E484K, N501Y), Gamma (K417T, E484K, N501Y), and Delta (L452R, T478K, P681R and occasionally E484Q) variants may lead to small changes in the structure of the S protein and RBD^8^ leading to a higher infection rate and reduction of acquired immunity by neutralizing antibodies. As previously reported, we observed a substantial reduction of neutralization activity in the plasma of convalescent patients collected at early and more particularly, late phase of convalescence against the Beta, Gamma and to a lower extend Delta^34^. The N501Y mutation in the Alpha variant RBD does not alter the neutralizing ability of plasma antibodies from naturally infected individuals^35^ while the Beta (B.1.351) and Gamma (P.1) carrying the N501Y and E484 mutations are more resistant to the neutralizing activity from convalescent and vaccine immune sera as well as neutralizing antibodies^7,11,36,37^. The substitution at position E484 in the RBD of pseudovirus or recombinant virus is known to confer resistance to neutralization by convalescent human sera^12,13^. The L452R mutation found in Delta was shown to impair neutralization by antibodies^38^ while the T478K mutation in the RBD is unique to Delta and close to the E484K mutation. Recent studies suggest that immunity from infection with prior lineage of the SARS-CoV-2 virus provides a lower protection against reinfection with Delta^39^, although it could still reduce disease severity and hospitalization^40^.

Memory B and T cell immune responses with SARS-CoV-2 have been detected in convalescent individuals but clear correlates for protective immunity have yet to be defined. RBD and S-specific memory B cells are very rare in unexposed individuals but start to appear within 2 weeks after SARS-CoV-2 infection^5,6,41^. RBD- and S-specific memory B cells steadily increased over the following months and were still present more than 6 months after initial infection^5,6^. To date, only one study has examined the RBD-specific memory B cell response up to 12 months after onset of symptoms^7^. Using ELISpot, we measured the number of B cells secreting IgG antibodies specific for SARS-CoV-2 RBD which are the most numerous and more likely to produce neutralizing antibodies. In accordance with our previous results^7^, we observed that while serum anti-RBD antibodies peaked 15-30 days post-infection and gradually decrease, RBD-specific memory B cells increased with time, reaching a maximum at 3-6 months post-infection, to a similar level induced in individuals receiving two doses of Pfizer vaccines. More importantly, the member B cell response persist without further significant decrease for up to 15 months post-symptom onset, irrespective of disease severity. This study highlights that a decline in serum antibodies in convalescent patients may not reflect waning immunity, but rather a contraction of the immune response, with the development and persistence of virus-specific, long-lived memory B cells in the bone marrow. It has been shown that previous infection with SARS-CoV-2 increases both the number of RBD binding memory cells and neutralizing antibody titers after a single dose of vaccine^7,37,42^ suggesting the induction of plasma cells differentiation from the memory B cell compartment^17^.

Early T cell responses during COVID-19 have been correlated with rapid viral clearance and reduced disease severity^43^. In accordance with previous results, memory IL-2 and/or IFN-γ−producing T cells specific to M, N nonstructural proteins, as well as S protein, were generated in the majority of convalescent patients following SARS-CoV-2 infection^43–46^. Functional SARS-CoV-2 specific T cells were detected at low level within 2 to 4 weeks from onset of symptoms and reached a peak between 3 and 6 months at levels similar to that of fully vaccinated individuals^4^. Specific IFN-γ producing Th1 CD4^+^ and cytotoxic CD8^+^ effector cells are considered important in supporting immunity against SARS-CoV-2^47^. Polyfunctional T cells, secreting more than one cytokine, are also typically associated with superior control of pathogens and may be important to prevent reinfection by SARS-CoV-2^48,49^. The presence of dual positive IFN-γ and IL-2 S1-specific T cells that preferentially retain characteristics of both effector function and proliferative potential *in vivo* is indicative of strong and sustained S-specific T-cell immunity. Similar to previous results^5^, a decrease in specific T cell response, and more particularly in dual positive IFN-γ and IL-2 S1- and S N M O-specific T cells, was observed after 6 months. Nevertheless, memory T cell were observed in the majority of the patients between 12-15 months and a longer follow-up period with more participants will be necessary to evaluate the sustainability of the T cell response beyond 15 months post-infection.

In conclusion, we observed that circulating memory B and T cells, and neutralizing antibodies are present in the majority of convalescent patients around 15 months after SARS-CoV-2 infection, demonstrative of a long-lasting immune response. Recent studies have shown that although the Pfizer-BioNTech and AstraZeneca vaccines were effective in reducing the risk of infection and COVID-19 hospitalization caused by Delta, these effects on infection appeared to be diminished when compared to those with the Alpha variant^40^. Our neutralization data suggest that immunity develops during earlier waves of infection may not be fully protective against reinfection with Delta and other VOCs indicating that convalescent patients may still benefit from vaccination. Furthermore, although two-doses of vaccines could trigger rapid and robust immune responses as observed in naturally infected individuals, the longevity of immunity in vaccinated individuals should be followed up in future studies.

## Limitation of studies

Potential limitation to our study includes the limited number of convalescent patients analyzed between 12 and 15 months. In addition, cell phenotyping was not performed, and it was not possible to distinguish whether the T cells measured at early time points were effector or memory cells. More individuals and cell phenotyping might help to characterize better the longevity of the immune response development, the persistence of memory B and T cells following natural infection, and the effect of disease severity.

## Acknowledgments

We thank all of the patients, blood donors, clinicians and nurses for their contributions. This project was supported by the European Union’s Horizon 2020 research and innovation programme (ATAC, No. 101003650), the Italian Ministry of Health (Ricerca Finalizzata grant no. GR-2013-02358399), the Center for Innovative Medicine and the Swedish Research Council. J.A. was supported by the SciLifeLab/ Knut and Alice Wallenberg national COVID-19 research program project grant 2020.

## Author contributions

F.Z. performed the ELISA experiment. A.P., L.D., I.C., E.P., J.C.S., J. Attevall, F. Bergami, M.C., and F. Baldanti contributed to patient data curation, sample preparation, and the neutralization assay. L.D., Y.W., and S.V. designed and/or performed the ELISpot and FluoroSpot experiments. M.V., M. Sambo, V.Z., E.A., R.B., T.O., and F.M. contributed to the enrollment of patients, patient management, and collection of clinical data. J. Andréll, F. Bertoglio, M. Schubert, and M.H. performed the production and purification of proteins from SARS-CoV-2 and variants. H.M., F.Z., L.D., I.C., H.W., M.K.-B., E.P., H.A., L.C., L.V, L.H., and Q.P.-H. contributed to the analysis and interpretation of data. H.M. and Q.P.-H. drafted the manuscript. H.M., A.P., Y.X., L.H., F.Baldanti, and Q.P.-H. conceived and supervised the study. All authors reviewed the manuscript.

## STAR Methods

### Resource Availability

#### Lead Contact

Further information and requests for resources and reagents should be directed to and will be fulfilled by the Lead Contact, Harold Marcotte.

#### Materials Availability

RBD and S1-S2 proteins can be generated and shared on a collaborative basis.

#### Data and Code Availability

All relevant data outputs are within the paper and its supplementation information.

## Experimental Model and Subject Details

### Study design and participants

Previously enrolled study participants were asked to return for a follow-up visit at the Fondazione IRCCS Policlinico San Matteo in Pavia, Italy and Karolinska Institutet, Stockholm, Sweden. Eligible participants were 18 years of age with a history of participation in prior study visit(s) of our longitudinal cohort study of COVID-19 recovered individuals^6^. All participants had a confirmed history of SARS-CoV-2 infection, who had tested PCR- or serology-positive for SARS-CoV-2^6^. Of those patients, 7 Italian and 4 Swedish patients returned for a follow-up sample between February 1 and June 10, 2021. In addition, 21 Italian and 28 Swedish convalescent patients were recruited between November 18 and June 16, 2021. Study inclusion criteria included subjects over 18 years of age, who were willing and able to provide informed consent, confirmed positivity of SARS-CoV-2 by real-time RT-PCR targeting the E and RdRp genes according to Corman *et al*.^50^ protocols and monitored until two subsequent samples with negative results.

Blood sample was taken from 53 patients at one single time point between day 15 and 452 after symptoms onset while 7 patients had blood taken at two time points for a total of 67 samples including 20 between 241 to 452 days (9-15 months). Results obtained from the new recruited and new follow-up samples were merged with previously published data assessing the immune response to SARS-CoV-2 up to 6-8 months^6^. Following merging, a total of 188 blood samples were collected from 136 patients, 98 Italians and 38 Swedish, for the entire 15-months follow-up (Figure 1). Ninety-four donors had blood drawn at one single time point ranging from 7 to 452 days after symptom onset while 35, 5, and 1 donors had blood taken at two, three and four time points, respectively. For detailed participant characteristics see Supplementary Tables S1).

Disease severity was defined as mild (non-hospitalized), moderate (hospitalized, with lower respiratory tract infection, with dyspnea or not, but without oxygen support), severe (infectious disease/sub intensive ward with a need for oxygen and/or positive chest computed tomography scan, severe lower tract infections, with any oxygen support) and critical (intensive care unit (ICU) patients, intubated or with extracorporeal membrane oxygenation procedures)^6^.

The demographic and clinical characteristics of the patients for the entire 15 months follow-up are detailed in Table S1 and summarized in Table S2. The Italian patients, 58 (59.2%) males and 40 (40.8%) females, had a median age of 66.0 years (range 22-89). The degree of clinical severity of COVID-19 in the cohort was mild (n = 8), moderate (n = 17), severe (n = 67) and critical (n = 6). The Swedish patients had a median age of 44 years (range 18-75) with 16 (42.1%) males and 22 (57.9%) females and all 38 of them had mild symptoms.

In addition, serum samples from 108 anonymized individuals (16 to 80 years of age), collected before the SARS-CoV-2 pandemic (1995 to 2005) were used as historical negative controls for the ELISA. PBMCs from four healthy controls (median age 41 years, range 39-50) and seven additional buffy coats collected in Sweden before the SARS-CoV-2 pandemic (2011-January 2020) were included as negative controls for the B and T cell assays. Patients and samples tested in different assays are summarized in a flow chart (Figure 1).

Furthermore, 23 individuals (9 males and 14 females) with a median age of 40 years (range 23-64) who received the mRNA Comirnaty vaccine (Pfizer-BioNTech) were included for comparison. Vaccinated individuals were sampled 14-35 days after the first dose and 14-36 days after the second dose. Seven individuals were samples after the first and second doses and 16 were sampled either after the first (n=8) or second (n=8) dose for a total of 15 samples after each dose (Figure 1, Table S3).

The study in Italy was performed under the approval of the Institutional Review Board of Policlinico San Matteo (protocol number P_20200029440). The study in Sweden was approved by the ethics committee in Stockholm (Dnr 2020-02646). All participants provided written informed consent before participation in the study.

## Method Details

### Production of SARS-CoV-2 RBD and S proteins

RBD-His protein was expressed in Expi293 cells and purified on Ni-NTA resin (#88221, Thermo Fisher) followed by size-exclusion chromatography on a Superdex 200 gel filtration column in PBS.^51^ S1-S2-His (referred as S) protein from Wuhan-Hu-1 strain and RBD from variants (Alpha, Beta, Gamma, and Delta) were expressed baculovirus-free in High Five insect cells^52^ and purified on HisTrap excel column (Cytiva) followed by preparative size exclusion chromatography on 16/600 Superdex 200 pg column (Cytiva)^53,54^.

### Detection of antibodies specific to SARS-CoV-2

Levels of anti-S and anti-RBD IgM, IgA and IgG antibodies were determined by ELISA as previously described^6^. High-binding Corning Half area plates (Corning #3690) were coated overnight at 4°C with purified S or RBD protein derived from wild-type virus (1.7 μg/ml for IgM and IgG; 2.0 μg/ml for IgA) in PBS. Serum or plasma diluted 1:3200 (S IgM), 1:6400 (S IgG), 1:1600 (S IgA; RBD IgM, IgA, IgG) in 0.1% BSA in PBS, was incubated for 1.5h at room temperature. Plates were then washed and incubated for 1h at room temperature with horseradish peroxidase (HRP)-conjugated goat anti-human IgM (Invitrogen #A18835), goat anti-human IgA (Jackson #109-036-011), or goat anti-human IgG (Invitrogen #A18805), (all diluted 1:15 000 in 0.1% BSA-PBS). Bound antibodies were detected using tetramethylbenzidine substrate (Sigma #T0440). The color reaction was stopped with 0.5M H_2_SO_4_ and absorbance was measured at 450nm. Antibody levels were presented as arbitrary units (AU/ml), based on a standard curve made from a serially diluted highly positive serum pool. In-house standards made by pooled highly positive serum were calibrated by using the WHO International Standard for anti-SARS-CoV-2 immunoglobulin (NIBSC, 20/136). 1 AU/ml of in-house serum standard equal to 30.20 AU/ml IgM, 185.59 AU/ml IgA and 4.10 AU/ml IgG of the WHO international standards, respectively. Cutoff values for antibody positivity were determined based on receiver operating characteristic curves with data from convalescent COVID-19 patients and negative historical control samples. The cutoff value for positivity was set at > 2.5 AU/ml for anti-S IgM, > 0.5 AU/ml for anti-S IgA, > 0.03 AU/ml for anti-S IgG, > 8.4 AU/ml for anti-RBD IgM, > 0.08 AU/ml for anti-RBD IgA, and > 14.8 AU/ml for anti-RBD IgG, giving a specificity of 96% for IgM, 99% for IgA and 97% IgG.

For assessing the anti-RBD IgG binding activity against VOCs, plates were coated with RBD derived from Alpha, Beta, Gamma, and Delta (1.7 μg/ml) and 1:1600 dilution of selected serum or plasma collected from February to May 2021 at early (15-106 days, median day of 24) and late (259-452 days, median day of 370) phase of convalescence were added. Horseradish peroxidase (HRP)-conjugated goat anti-human IgG and tetramethylbenzidine substrate were added as described above and the color reaction was stopped with 0.5M H_2_SO_4_ after 10 min incubation. All samples were run in the same experiment and the results expressed as OD_450_. As an internal control, a diluted in-house developed monoclonal antibody cross-binding all RBD variants was used.

### Neutralization

SARS-CoV-2 strain G614 and VOCs (Alpha, Beta, Gamma, and Delta) were isolated from patients in Pavia, Italy and identified through next-generation sequencing method. A previously described microneutralization assay was used to determine the titers of neutralizing antibodies against SARS-CoV-2 strains in 171 samples^55,56^. The neutralizing titer (NT_90_) was the maximum dilution giving a reduction of 90% of the cytopathic effect. The cutoff for positivity was ≥1:10.

### Isolation of PBMCs and RNA

PBMCs were isolated from blood or buffy coat samples by standard density gradient centrifugation using Lymphoprep (Axis-Shield) and were cryopreserved and stored in liquid nitrogen until analysis. Total RNA was extracted from PBMCs by using RNeasy mini kit according to the manufacturer’s protocol (Qiagen).

### ELISPOT and FluoroSpot

PBMCs were incubated for four days in RPMI-1640 medium with 10% FCS, supplemented with the TLR7 and TLR8 agonist imidazoquinoline resiquimod (R848, 1 μg/ml; Mabtech AB, Nacka, Sweden), and recombinant human IL-2 (10 ng/ml) for stimulation of memory B cells. The number of B cells secreting SARS-CoV-2 RBD-specific IgG and total IgG were measured using the Human IgG SARS-CoV-2 RBD ELISpot^PLUS^ kit (Mabtech AB)^6^.

S1 and S N M O specific IFN-γ and IL-2-secreting T cells were detected using the Human IFN-γ/IL-2 SARS-CoV-2 FluoroSpot^PLUS^ kit (Mabtech AB)^6^. The plates pre-coated with capturing monoclonal anti-IFN-γ and anti-IL-2 were incubated overnight in RPMI-1640 medium containing 10% FCS supplemented with a mixture containing the SARS-CoV-2 peptide pool (scanning or defined pools), anti-CD28 (100 ng/ml) and 300 000 cells per well in humidified incubators (5% CO_2,_ 37°C). The SARS-CoV-2 S1 scanning pool contains 166 peptides from the human SARS-CoV-2 virus (#3629-1, Mabtech AB). The peptides are 15-mers overlapping with 11 amino acids, covering the S1 domain of the S protein (amino acid 13-685). The SARS-CoV-2 S N M O defined peptide pool contains 47 synthetic peptides binding to human HLA, derived from the S, N, M ORF3a and ORF7a proteins (#3622-1, Mabtech AB)^57^. The SARS-CoV-2 S2 N defined peptide pool contains 41 synthetic peptides binding to human HLA derived from the S and N proteins of the SARS-CoV-2 virus (#3620-1, Mabtech AB)^58^.

Results of ELISpot and FluoroSpot assays were evaluated using an IRIS-reader and analyzed by the IRIS software version 1.1.9 (Mabtech AB). The results were expressed as the number of spots per 300 000 seeded cells after subtracting the background spots of the negative control. The cutoff value was set at the highest number of specific B- and T cell spots from the negative controls. The number of SARS-CoV-2 specific T cells (per 300 000 cells) producing either IL-2, IFN-γ or both IL-2 and IFN-γ (IL-2/IFN-γ) were plotted.

### Quantification and Statistical analysis

Mann-Whitney U test was used for comparisons between groups in anti-SARS-CoV-2 antibody levels, neutralization titers and numbers of specific memory B and T cells. Correlation analysis was performed using Spearman’s rank correlation. The number of patients showing neutralization activity against variants was compared by Fisher’s exact test. A Wilcoxon signed-rank test was used for comparison of paired samples. Continuous decay (linear regression), one-phase decay, two-phase decay, polynomial second order or log-normal of non-transformed and log_2_ transformed data were assessed with the best fitting statistical model chosen based on the F test. The half-lives of antibody and neutralization titers were estimated by a one-phase exponential decay model using non-transformed and log_2_ transformed data, respectively. All analyses and data plotting were performed using GraphPad or R version 3.6.1.

## Supplementary excel table title and legend

**Table S1** Demographic and clinical characteristics of the COVID-19 patients. Related to Figure 1.

## Supplemental information

**Figure S1. Antibody and neutralization titers in relation to severity. Related to Figures 2-4**. Levels of anti-RBD, anti-S and neutralization activity according to severity using cross-sectional (all samples) (A-C) and longitudinal (paired samples) (D-F) analysis. The antibody titers and NT_90_ decay curves (in black) together with corresponding half-lives (t_1/2_) were estimated by a one-phase decay model. Symbols represent individual subjects. The dashed red line indicates the cutoff value.

**Figure S2. Correlation between the level of anti-SARS-CoV-2 antibodies and neutralizing antibody titers. Related to Figures 2 and 4**. Levels of anti-RBD (A) and anti-S (B) IgM, IgA, and IgG antibodies significantly correlated with neutralization antibody titers measured against SARS-CoV-2 virus in COVID-19 patient samples (n=168 for each antibody isotype). Symbols represent individual subjects. Spearman’s rank correlation. Significant if p < 0.05.

**Figure S3. Cross-sectional analysis of SARS-CoV-2-specific memory B cell responses in COVID-19 patients in relation to disease severity. Related to Figure 5**. Dynamics of RBD-specific memory B cells levels in samples from COVID-19 patients with mild/moderate (A) and severe/critical (B) symptoms with the corresponding log-normal fitting curve (in black). The results were expressed as the number of spots per 300,000 seeded cells after subtracting the background spots of the negative control. Symbols represent individual subjects. The cutoff value (dashed red line) was set at the highest number of specific B cell spots for the negative controls (> 12 spots / 300,000 cells).

**Figure S4. Cross-sectional and longitudinal analysis of S N M O-specific memory T cell responses in COVID-19 patients. Related to Figure 6**. (A-C) Dynamics of S N M O-specific memory IL-2, IFN-γ, and IL-2/IFN-γ-producing T cells with the corresponding second order polynomial fitting curve (in black). (D-F) T cells were measured in control (n =11), COVID-19 samples at 5 study periods: 14-30 days (n = 11), 31-90 (n = 9), 91-180 days (n = 19), 181-365 (n = 28), and 366-452 (n = 6) days after symptom onset, as well as vaccinee samples after first (n = 15) and second (n = 11) dose. For longitudinal analysis, samples were taken at two (n = 10) or more (n = 5) time points, and further comparisons were made between paired samples (n = 8) at two time points ranging from 6 to 15 months (TP1: 181-300 and TP2: 301-452 days after symptoms onset; right panel). The results were expressed as the number of spots per 300,000 seeded cells after subtracting the background spots of the negative control. The horizontal black lines indicate the median value and 95% CI of the group (D-I). The cutoff value (dashed red line) was set at the highest number of specific T cell spots for the negative controls (> 6 to 13 spots / 300 000 seeded cells depending of the T cell population). Mann-Whitney U test. **p ≤ 0.01, ***p ≤ 0.001, and ****p ≤ 0.0001.

**Figure S5. Cross-sectional analysis of SARS-CoV-2-specific memory T cell responses in COVID-19 patients in relation to disease severity. Related to Figures 6 and S4**. Dynamics of S1 (A-C) and S N M O (D-F) peptide pools-specific memory IL-2, IFN-γ, and IL-2/IFN-γ-producing T cells in samples from COVID-19 patients with mild/moderate (A) and severe/critical (B) symptoms with the corresponding second order polynomial (mild/moderate) or log-normal (severe/critical) fitting curve (in black). The results were expressed as the number of spots per 300,000 seeded cells after subtracting the background spots of the negative control. Symbols represent individual subjects. The cutoff value (dashed red line) was set at the highest number of specific T cell spots for the negative controls (> 6 to 13 spots / 300 000 seeded cells depending on the T cell population). Mann-Whitney U test.

**Figure S6. Comparison of SARS-CoV-2-specific memory B and T cells in COVID-19 patients according to longevity and severity. Related to Figures 5, 6, and S4**. Number of B cells (A), and S1 (B) and S N M O (C) peptide pools - specific T cells in mild/moderate and severe/critical COVID-19 patients between 6 and 15 months after symptoms onset. Symbols represent individual subjects. The horizontal black lines indicate the median value and 95% CI of the group. Mann-Whitney U test.

**Figure S7. Heat-map representation of the SARS-CoV-2 specific adaptive immune responses. Related to Figures 2, 4-6 and S4**. For subjects with available data on all three “arms” of adaptive immunity (serum anti-RBD IgM, IgA, IgG and neutralization titers (NT90), the number of RBD-specific memory B cells, and the number of T cells specific for the virus protein-derived peptides pools producing IFN-γ, IL-2, or IFN-γ and IL-2 (Dual), are indicated. For each arm of immunity, the relative intensity of signals varies from no (grey), low (yellow) to high (dark green) signal. The gender (male: blue, female: red), age (from 22 (white) to 86 (dark pink) years), severity (from mild (light orange) to critical (dark orange)), number of signals (1, 2 or 3), and the number of days after symptoms onset are shown.

**Table S2. Summary of demographic and clinical characteristics in convalescent patients. Related to Figure 1**.

**Table S3. Characteristics of the cohort of vaccinated recipients sampled after the first and second dose. Related to Figure 1**.

