## Supplemental data for "Immunity to SARS-CoV-2 up to 15 months after infection"

**Table of contents**

**Figure S1.** Antibody and neutralization titers in relation to severity. Related to Figures 2-4.

**Figure S2.** Correlation between the level of anti-SARS-CoV-2 antibodies and neutralizing antibody titers. Related to Figures 2 and 4.

**Figure S3.** Cross-sectional analysis of SARS-CoV-2-specific memory B cell responses in COVID-19 patients in relation to disease severity. Related to Figure 5.

**Figure S4.** Cross-sectional and longitudinal analysis of S N M O-specific memory T cell responses in COVID-19 patients. Related to Figure 6.

**Figure S5.** Cross-sectional analysis of SARS-CoV-2-specific memory T cell responses in COVID-19 patients in relation to disease severity. Related to Figures 6 and S4.

**Figure S6.** Comparison of SARS-CoV-2-specific memory B and T cells in COVID-19 patients according to longevity and severity. Related to Figures 5, 6, and S4.

**Figure S7.** Heat-map representation of the SARS-CoV-2 specific adaptive immune responses. Related to Figures 2, 4-6 and S4.

**Table S2.** Summary of demographic and clinical characteristics in convalescent patients. Related to Figure 1.

**Table S3.** Characteristics of the cohort of vaccinated recipients sampled after the first and second dose. Related to Figure 1.

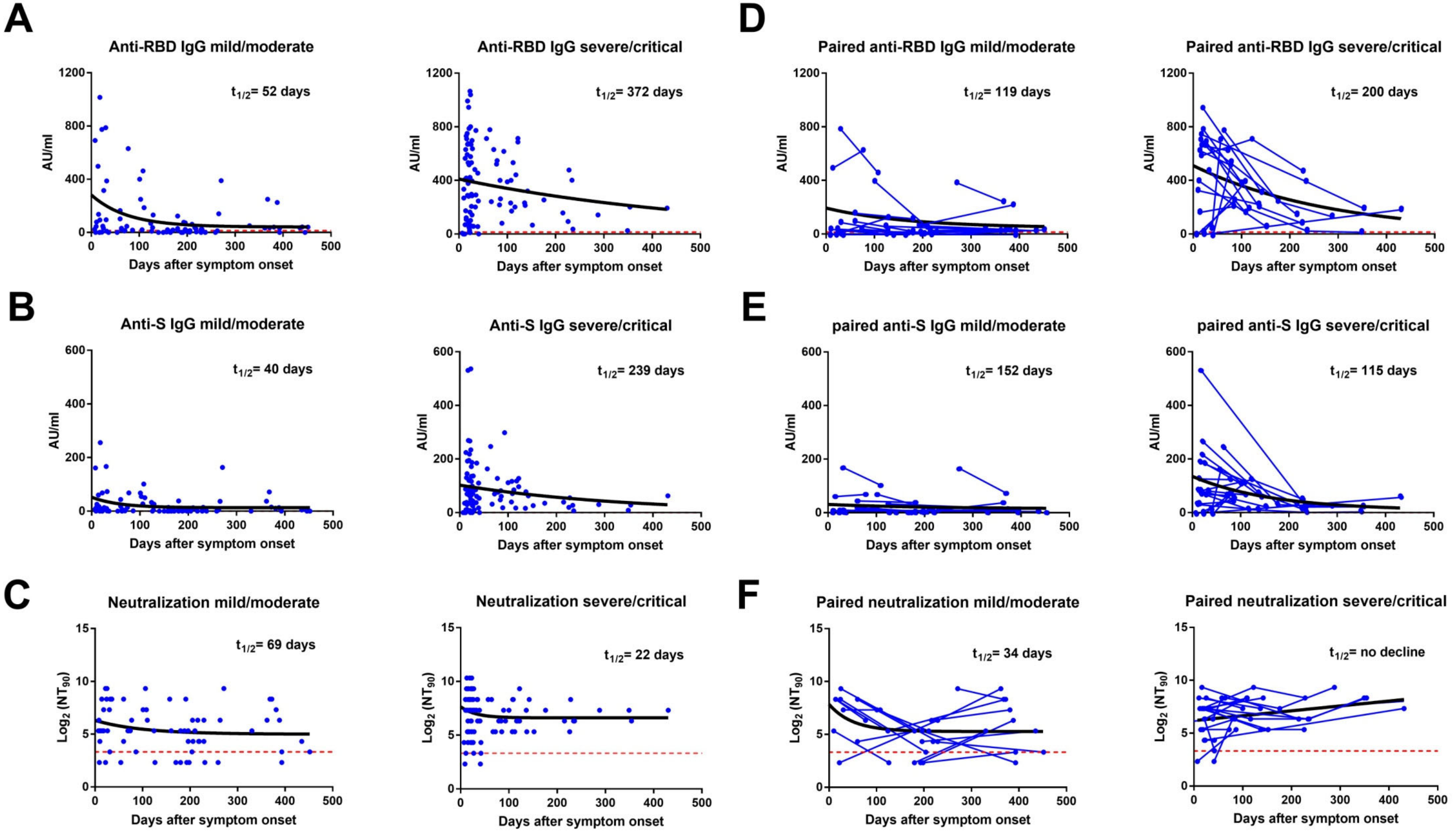


**Figure S1. Antibody and neutralization titers in relation to severity. Related to Figures 2-4.** Levels of anti-RBD, anti-S and neutralization activity according to severity using cross-sectional (all samples) (A-C) and longitudinal (paired samples) (D-F) analysis. The antibody titers and $NT_{90}$ decay curves (in black) together with corresponding half-lives ($t_{1/2}$) were estimated by a one-phase decay model. Symbols represent individual subjects. The dashed red line indicates the cutoff value.

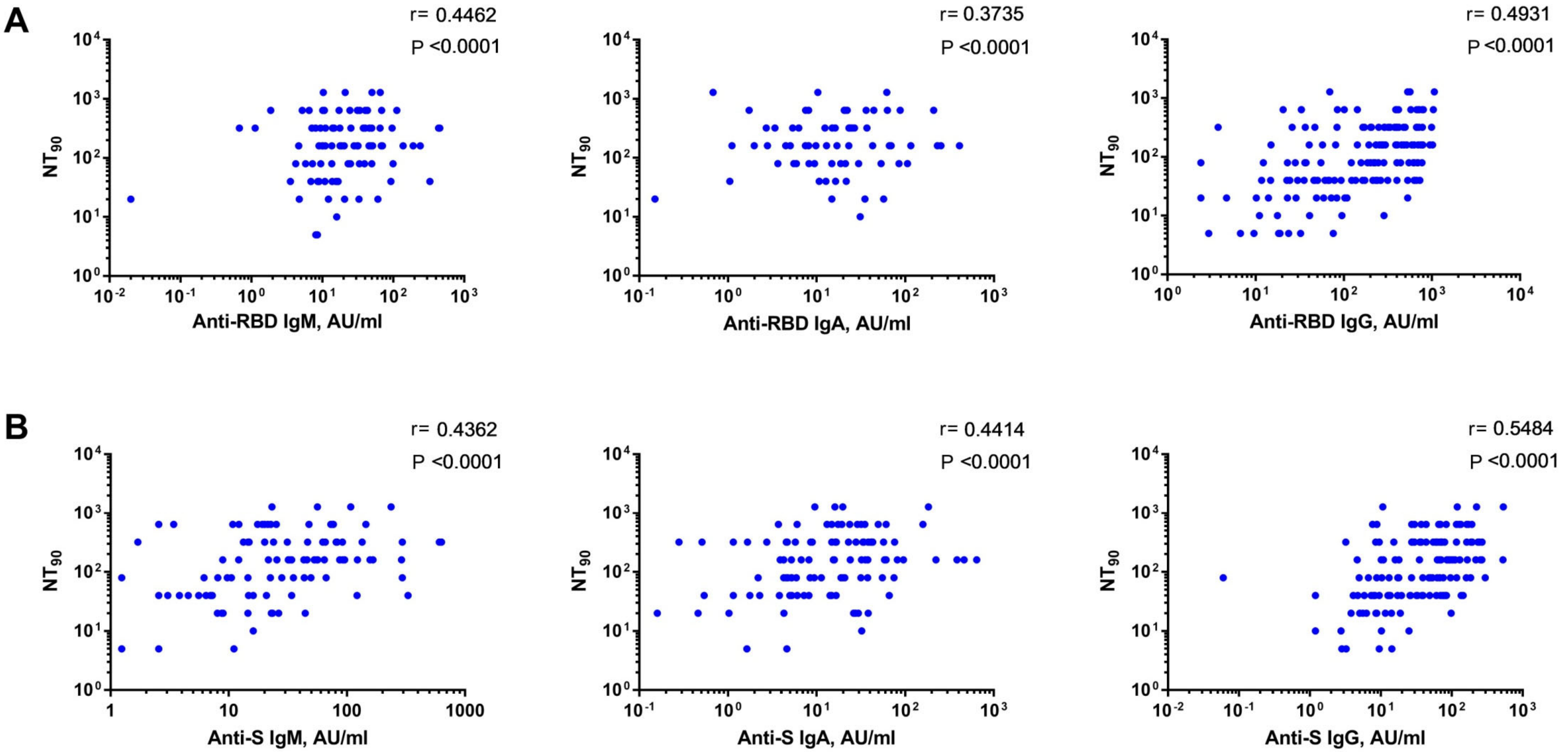


**Figure S2. Correlation between the level of anti-SARS-CoV-2 antibodies and neutralizing antibody titers. Related to Figures 2 and 4.** Levels of anti-RBD (A) and anti-S (B) IgM, IgA, and IgG antibodies significantly correlated with neutralization antibody titers measured against SARS-CoV-2 virus in COVID-19 patient samples (n=168 for each antibody isotype). Symbols represent individual subjects. Spearman's rank correlation. Significant if $p < 0.05$.

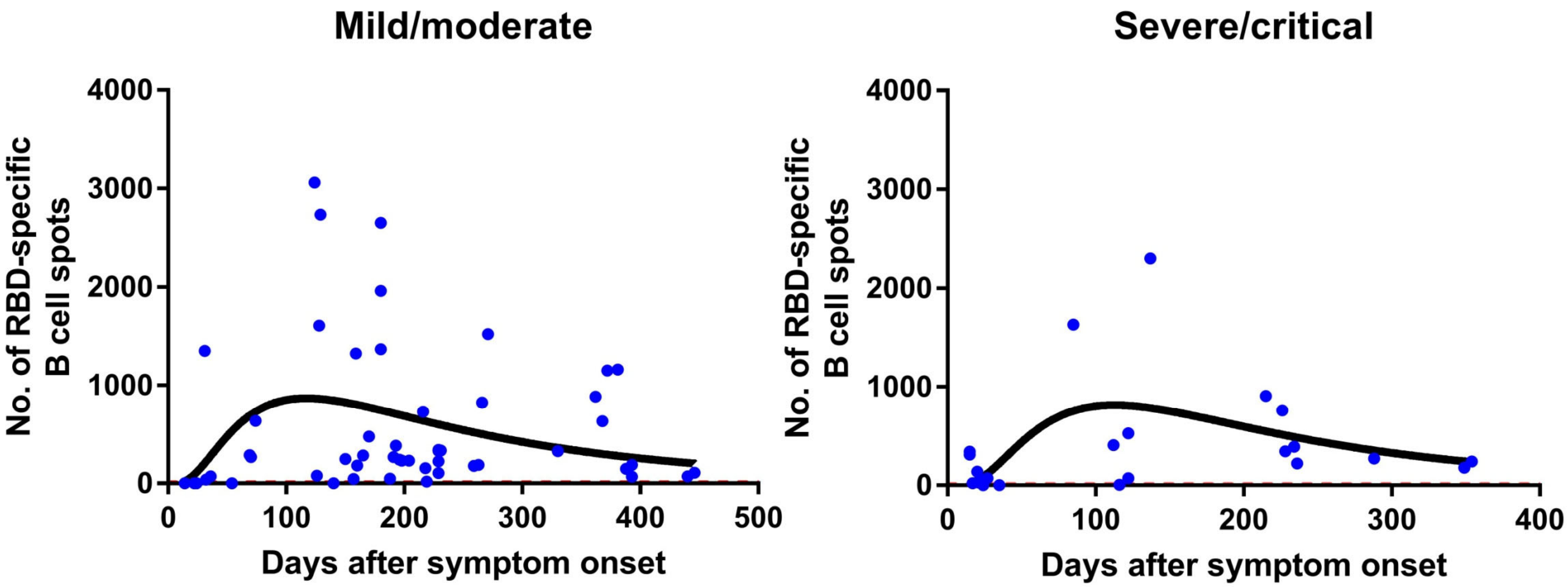


**Figure S3. Cross-sectional analysis of SARS-CoV-2-specific memory B cell responses in COVID-19 patients in relation to disease severity. Related to Figure 5.** Dynamics of RBD-specific memory B cells levels in samples from COVID-19 patients with mild/moderate (A) and severe/critical (B) symptoms with the corresponding log-normal fitting curve (in black). The results were expressed as the number of spots per 300,000 seeded cells after subtracting the background spots of the negative control. Symbols represent individual subjects. The cutoff value (dashed red line) was set at the highest number of specific B cell spots for the negative controls (> 12 spots / 300,000 cells).

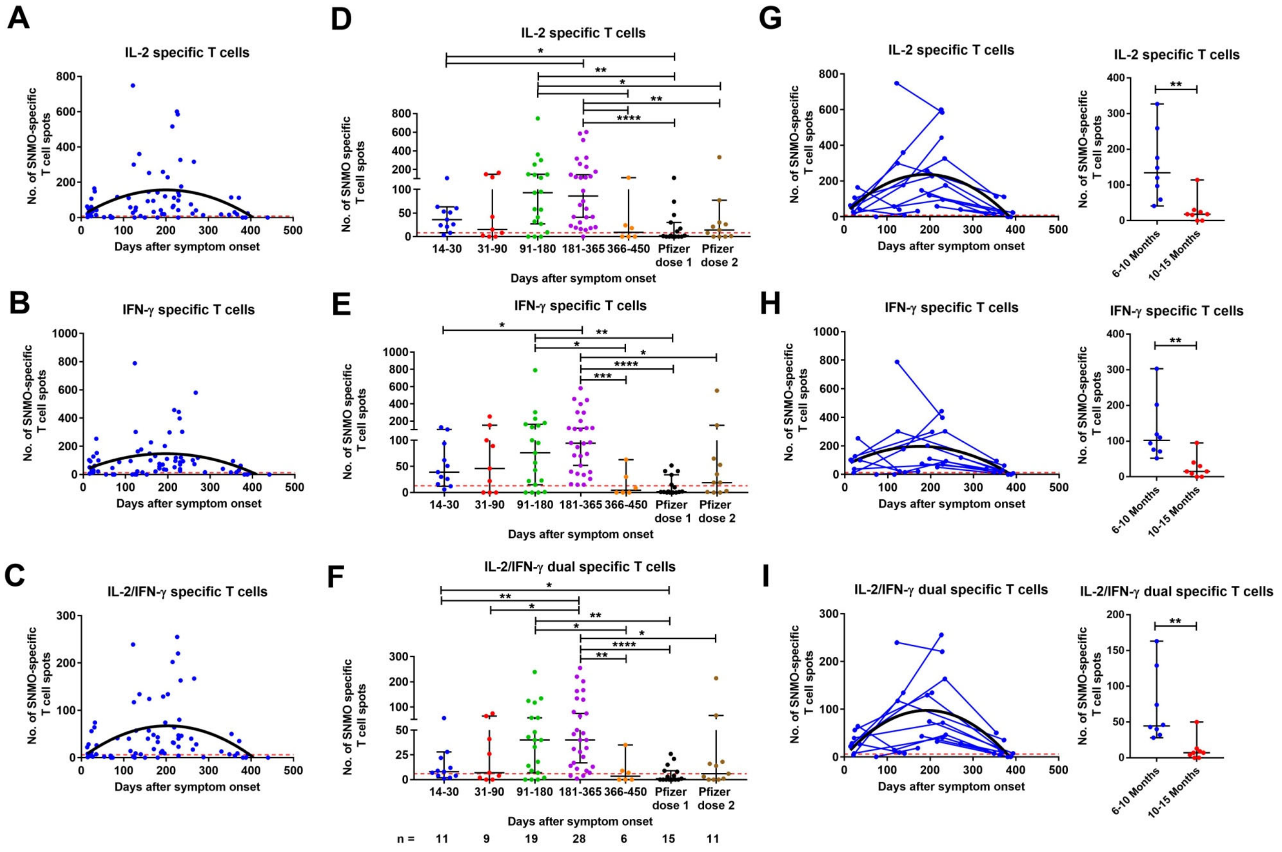


**Figure S4. Cross-sectional and longitudinal analysis of S N M O-specific memory T cell responses in COVID-19 patients. Related to Figure 6.** (A-C) Dynamics of S N M O-specific memory IL-2, IFN-γ, and IL-2/IFN-γ-producing T cells with the corresponding second order polynomial fitting curve (in black). (D-F) T cells were measured in control (n =11), COVID-19 samples at 5 study periods: 14-30 days (n = 11), 31-90 (n = 9), 91-180 days (n = 19), 181-365 (n = 28), and 366-452 (n = 6) days after symptom onset, as well as vaccinee samples after first (n = 15) and second (n = 11) dose. For longitudinal analysis, samples were taken at two (n = 10) or more (n = 5) time points, and further comparisons were made between paired samples (n = 8) at two time points ranging from 6 to 15 months (TP1: 181-300 and TP2: 301-452 days after symptoms onset; right panel). The results were expressed as the number of spots per 300,000 seeded cells after subtracting the background spots of the negative control. The horizontal black lines indicate the median value and 95% CI of the group (D-I). The cutoff value (dashed red line) was set at the highest number of specific T cell spots for the negative controls (> 6 to 13 spots / 300 000 seeded cells depending of the T cell population). Mann-Whitney U test. **p ≤ 0.01, ***p ≤ 0.001, and ****p ≤ 0.0001.

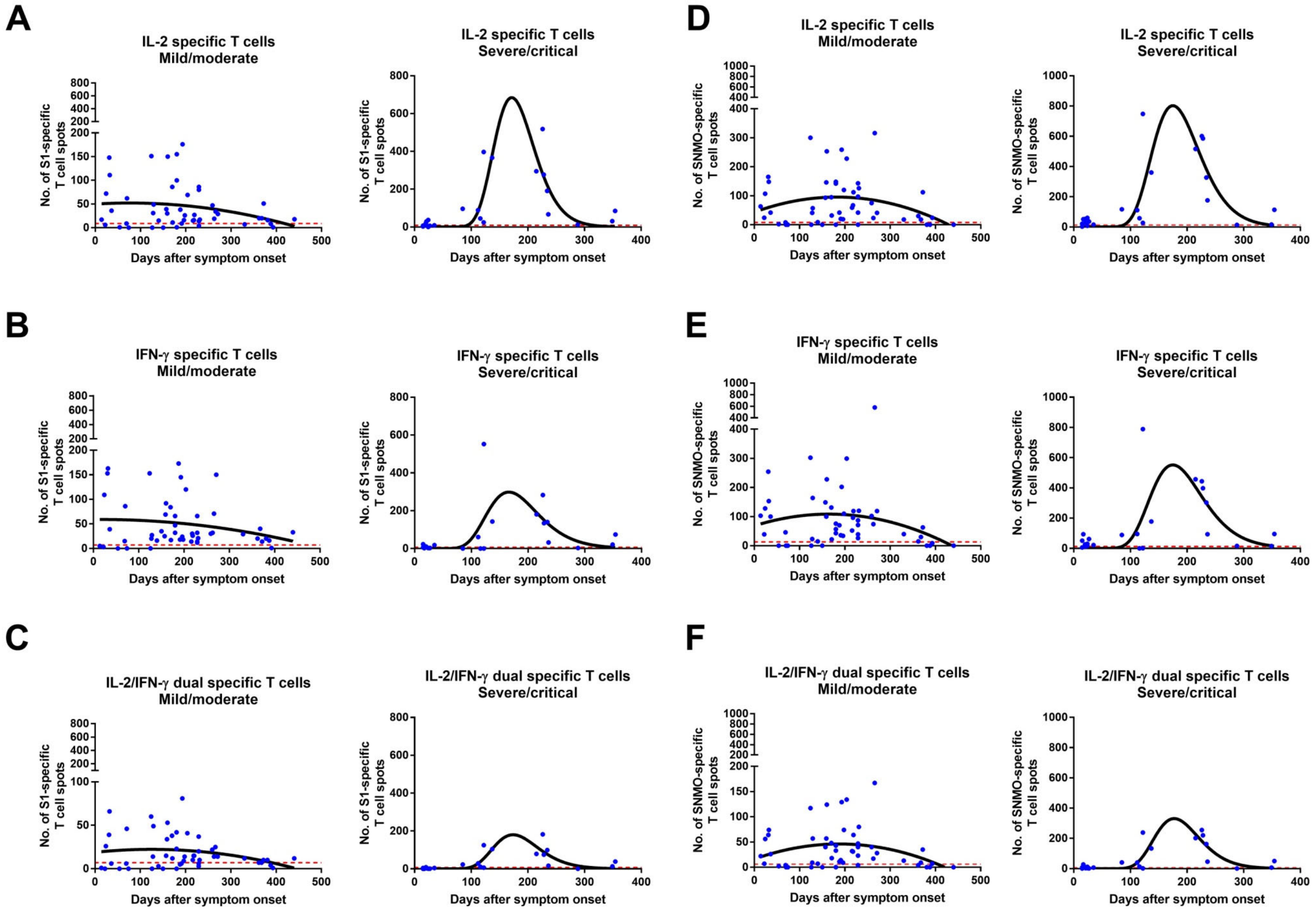


**Figure S5. Cross-sectional analysis of SARS-CoV-2-specific memory T cell responses in COVID-19 patients in relation to disease severity. Related to Figures 6 and S4.** Dynamics of S1 (A-C) and S N M O (D-F) peptide pools-specific memory IL-2, IFN-γ, and IL-2/IFN-γ-producing T cells in samples from COVID-19 patients with mild/moderate (A) and severe/critical (B) symptoms with the corresponding second order polynomial (mild/moderate) or log-normal (severe/critical) fitting curve (in black). The results were expressed as the number of spots per 300,000 seeded cells after subtracting the background spots of the negative control. Symbols represent individual subjects. The cutoff value (dashed red line) was set at the highest number of specific T cell spots for the negative controls (> 6 to 13 spots / 300 000 seeded cells depending on the T cell population). Mann-Whitney U test.

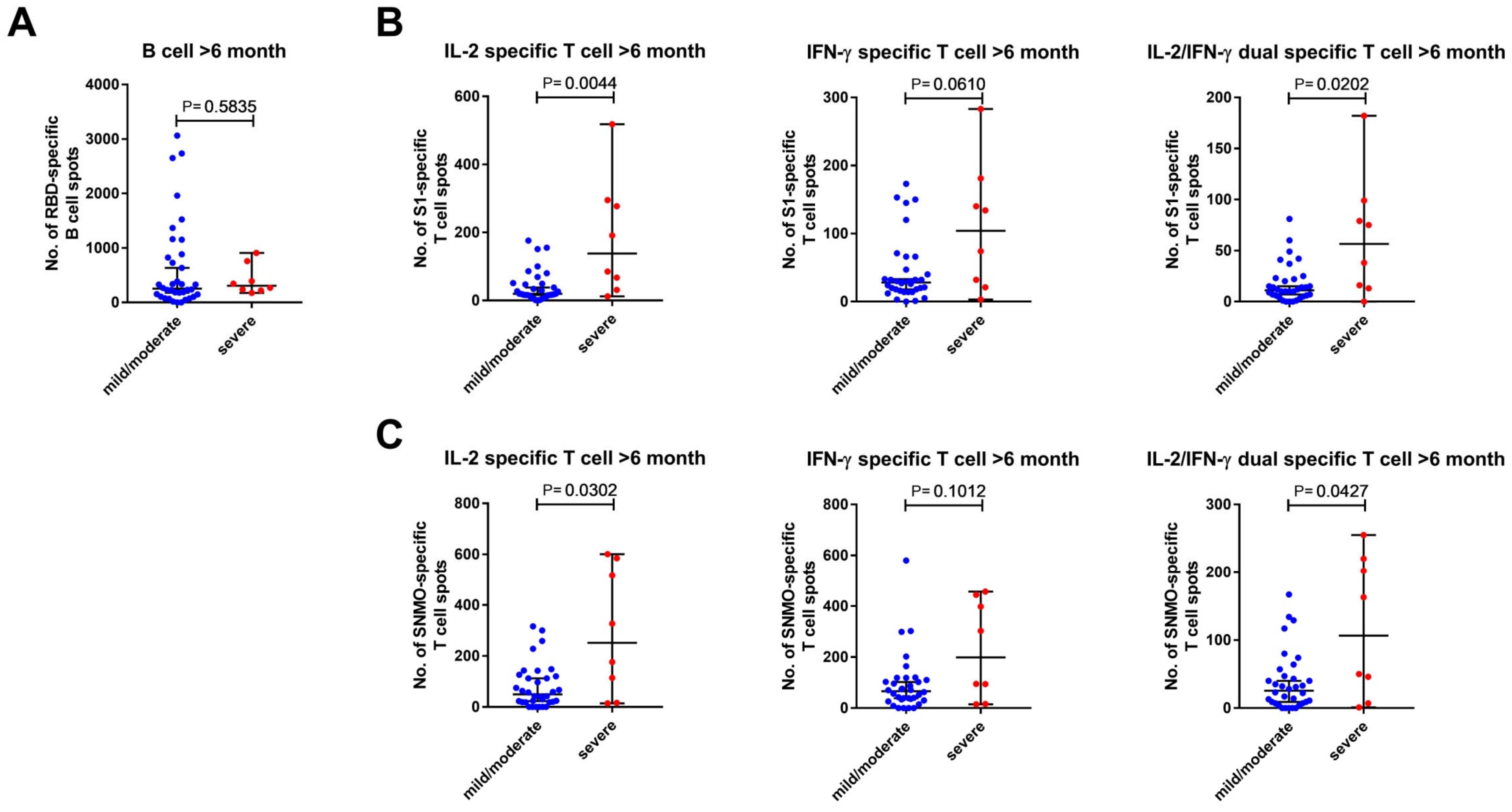


**Figure S6. Comparison of SARS-CoV-2-specific memory B and T cells in COVID-19 patients according to longevity and severity. Related to Figures 5, 6, and S4.** Number of B cells (A), and S1 (B) and S N M O (C) peptide pools - specific T cells in mild/moderate and severe/critical COVID-19 patients between 6 and 15 months after symptoms onset. Symbols represent individual subjects. The horizontal black lines indicate the median value and 95% CI of the group. Mann-Whitney U test.

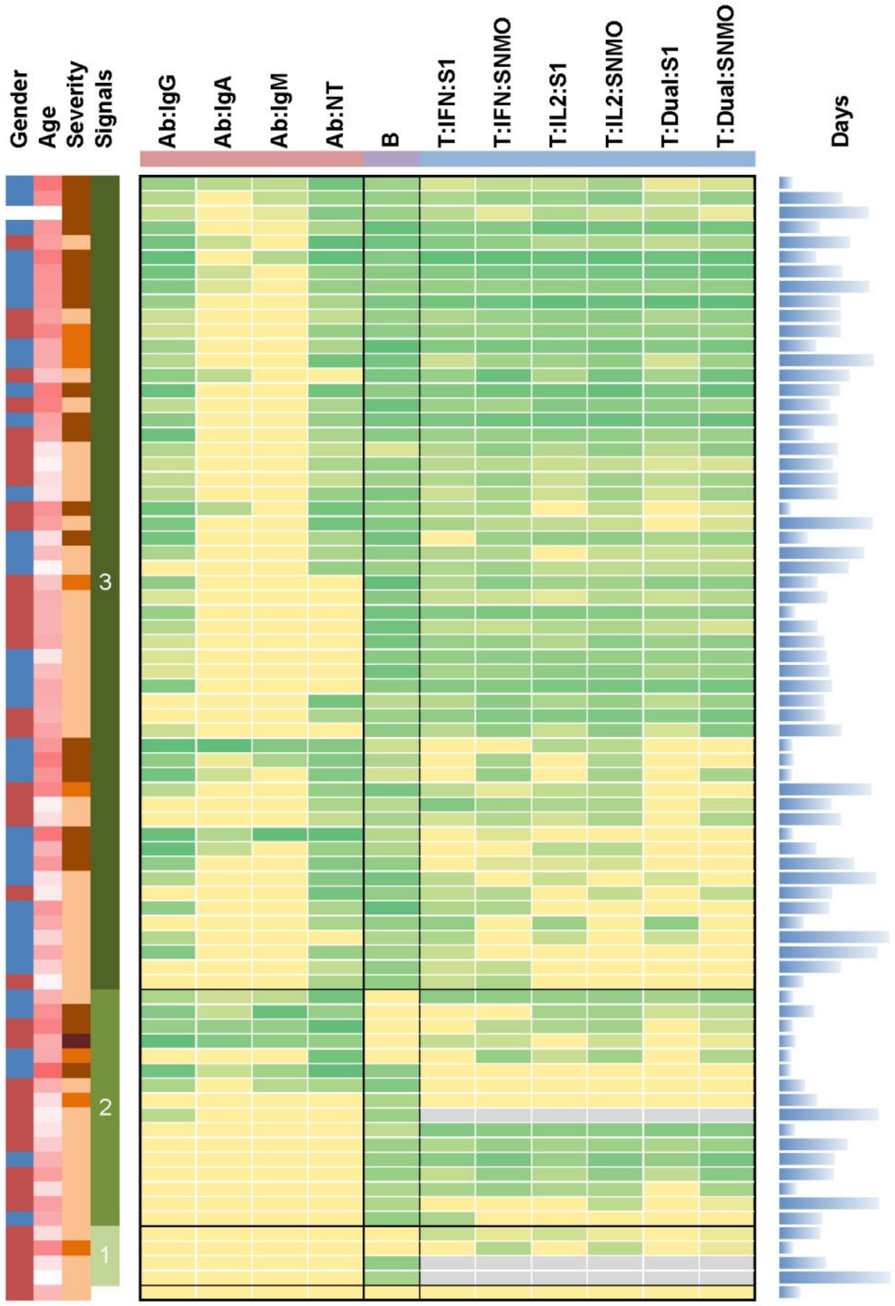


**Figure S7. Heat-map representation of the SARS-CoV-2 specific adaptive immune responses. Related to Figures 2, 4-6 and S4.** For subjects with available data on all three "arms" of adaptive immunity (serum anti-RBD IgM, IgA, IgG and neutralization titers (NT90), the number of RBD-specific memory B cells, and the number of T cells specific for the virus protein-derived peptides pools producing IFN-$\gamma$, IL-2, or IFN-$\gamma$ and IL-2 (Dual), are indicated. For each arm of immunity, the relative intensity of signals varies from no (grey), low (yellow) to high (dark green) signal. The gender (male: blue, female: red), age (from 22 (white) to 86 (dark pink) years), severity (from mild (light orange) to critical (dark orange)), number of signals (1, 2 or 3), and the number of days after symptoms onset are shown.

**Table S2. Summary of demographic and clinical characteristics in convalescent patients. Related to Figure 1.**

| | **Italian cohort** | **Swedish cohort** |
|---|---|---|
| No. | 98 | 38 |
| Age (Median, range), year | 66 (22-89) | 44 (18-75) |
| Male, % | 58 (59.2%) | 16 (42.1%) |
| Female, % | 40 (40.8%) | 22 (57.9%) |
| Disease severity, % | | |
| Mild | 8 (8.2%) | 38 (100%) |
| Moderate | 17 (17.3%) | 0 |
| Severe | 67 (68.4%) | 0 |
| Critical | 6 (6.1%) | 0 |

**Table S3. Characteristics of the cohort of vaccinated recipients sampled after the first and second dose. Related to Figure 1.**

| | 1 dose | 2 doses |
|---|---|---|
| No. of samples | 15[a] | 15 [a] |
| Age (Median, range) | 38 (23-62) | 41 (25-64) |
| Male, % | 9 (47%) | 8 (40%) |
| Female, % | 10 (53%) | 12 (60%) |

[a]Seven individuals were sampled after the first and second dose and 16 were sampled either after the first (n=8) or second (n=8) dose for a total of 15 samples after each dose.
