## Supplemental table S1 for "Immunity to SARS-CoV-2 up to 15 months after infection"

Table S1 Demographic and clinical characteristics of the COVID-19 patients. Related to Figure 1.

| New ID | score (mild, moderate, severe, critical) | Gender | Age | Cohort | days after onset of symptomes | symptoms | outcome | comorbidity | tumour | heart diseases | hypertension | diabetes | lung diseases | hcv | obesity | immunodeficiency | other comorb. |
|---|---|---|---|---|---|---|---|---|---|---|---|---|---|---|---|---|---|
| CP01 | severe | M | 72 | Italian | 13 | fever, dyspnea | alive | | no | yes | yes | no | no | no | yes | no | no |
| CP02 | severe | F | 78 | Italian | 18 | fever, cough | alive | | no | yes | yes | no | yes | no | no | no | no |
| CP03a | severe | M | 68 | Italian | 27 | fever, cough | alive | | no | no | no | no | no | no | no | no | no |
| CP03b | severe | M | 68 | Italian | 116 | | | | | | | | | | | | |
| CP03c | severe | M | 68 | Italian | 236 | | | | | | | | | | | | |
| CP03d | severe | M | 68 | Italian | 349 | | | | | | | | | | | | |
| CP04 | severe | M | 73 | Italian | 23 | fever, cough, dyspnea | death | | no | no | no | yes | no | yes | no | no | yes |
| CP06a | severe | M | 65 | Italian | 21 | fever, cough, dyspena, asthenia | alive | | no | no | no | yes | no | no | no | no | no |
| CP06b | severe | M | 65 | Italian | 137 | | | | | | | | | | | | |
| CP06c | severe | M | 65 | Italian | 226 | | | | | | | | | | | | |
| CP07a | mild | F | 61 | Italian | 25 | fever, cough | alive | | na | na | na | na | na | na | na | na | na |
| CP07b | mild | F | 61 | Italian | 229 | | | | | | | | | | | | |
| CP08a | moderate | F | 72 | Italian | 22 | fever, cough, dyarrhea | alive | | no | no | yes | no | no | no | no | no | no |
| CP08b | moderate | F | 72 | Italian | 229 | | | | | | | | | | | | |
| CP08c | moderate | F | 72 | Italian | 362 | | | | | | | | | | | | |
| CP09a | moderate | M | 56 | Italian | 14 | fever | alive | | no | no | yes | no | no | no | no | no | no |
| CP09b | moderate | M | 56 | Italian | 124 | | | | | | | | | | | | |
| CP09c | moderate | M | 56 | Italian | 372 | | | | | | | | | | | | |
| CP10a | moderate | F | 32 | Italian | 10 | fever, cough, pain | alive | | no | no | no | no | no | no | no | no | no |
| CP10b | moderate | F | 32 | Italian | 126 | | | | | | | | | | | | |
| CP11a | moderate | F | 44 | Italian | 9 | fever, dyarrhea | alive | | no | no | no | no | no | yes | no | no | no |
| CP11b | moderate | F | 44 | Italian | 129 | | | | | | | | | | | | |
| CP12 | severe | M | 70 | Italian | 23 | fever, cough, dyspena | alive | | no | no | no | yes | no | no | yes | no | no |
| CP13a | moderate | F | 70 | Italian | 30 | cough | alive | | no | yes | yes | no | no | no | no | no | no |
| CP13b | moderate | F | 70 | Italian | 108 | | | | | | | | | | | | |
| CP14 | severe | M | 78 | Italian | 12 | fever,asthenia | death | | yes | no | yes | yes | no | no | no | no | yes |
| CP15 | moderate | F | 86 | Italian | 18 | fever, cough | alive | | no | yes | yes | yes | no | no | no | no | no |
| CP16a | severe | F | 50 | Italian | 25 | fever, dyspnea, | alive | | no | no | yes | no | no | no | no | no | no |
| CP16b | severe | F | 50 | Italian | 100 | | | | | | | | | | | | |
| CP17a | severe | M | 42 | Italian | 10 | fever, cough, dyspnea | alive | | no | no | no | no | no | no | no | no | no |
| CP17b | severe | M | 42 | Italian | 136 | | | | | | | | | | | | |
| CP18 | severe | M | 53 | Italian | 18 | fever, dispnea | alive | | no | no | no | no | no | no | no | no | no |
| CP19a | severe | M | 77 | Italian | 17 | fever, cough | alive | | no | yes | no | no | no | no | no | no | yes |
| CP19b | severe | M | 77 | Italian | 122 | | | | | | | | | | | | |
| CP19c | severe | M | 77 | Italian | 228 | | | | | | | | | | | | |
| CP20 | severe | M | 57 | Italian | 20 | fever, cough, dyspnea | alive | | no | no | no | no | no | no | no | no | no |
| CP21 | severe | M | 81 | Italian | 28 | fever, cough | alive | | no | no | yes | no | no | no | no | no | no |
| CP23 | severe | M | 49 | Italian | 15 | fever, dyspnea | alive | | no | no | no | no | no | no | no | no | no |
| CP24 | severe | F | 60 | Italian | 39 | fever, dyspnea | alive | | no | no | yes | no | no | no | no | no | no |
| CP25 | moderate | F | 62 | Italian | 35 | fever, diarrhea | alive | | no | no | yes | no | no | no | no | no | yes |
| CP26 | critical | F | 55 | Italian | 35 | fever, cough | alive | | no | no | no | no | no | no | yes | no | no |
| CP27 | severe | F | 72 | Italian | 26 | fever, cough, dyspnea | alive | | yes | no | yes | no | no | yes | no | no | yes |
| CP28 | severe | F | 50 | Italian | 30 | fever, cough | alive | | no | no | yes | no | no | no | yes | no | yes |
| CP29 | critical | F | 53 | Italian | 23 | fever, dyspnea | alive | | no | no | yes | no | no | yes | no | no | no |
| CP29b | critical | F | 53 | Italian | 430 | | | | | | | | | | | | |
| CP30 | severe | M | 72 | Italian | 27 | fever, dyspnea | alive | | no | no | yes | yes | no | no | no | no | yes |
| CP31a | mild | F | 75 | Swedish | 60 | na | alive | | no | na | na | na | na | na | na | na | na |
| CP31b | mild | F | 75 | Swedish | 180 | | | | | | | | | | | | |
| CP33 | severe | M | 45 | Italian | 19 | fever,cough | alive | | no | no | yes | no | yes | no | no | no | no |
| CP34a | severe | M | 67 | Italian | 17 | fever | alive | | no | yes | yes | no | no | no | no | no | yes |
| CP34b | severe | M | 67 | Italian | 234 | | | | | | | | | | | | |
| CP34c | severe | M | 67 | Italian | 354 | | | | | | | | | | | | |
| CP35a | severe | M | 71 | Italian | 12 | fever, cough, dyspnea | alive | | no | no | yes | no | no | no | no | no | no |
| CP35b | severe | M | 71 | Italian | 152 | | | | | | | | | | | | |
| CP36 | moderate | M | 84 | Italian | 8 | fever, cough, dyspnea | alive | | no | no | yes | no | no | no | no | no | no |
| CP37a | severe | F | 43 | Italian | 20 | fever, cough, dyspnea | alive | | no | no | yes | yes | no | no | yes | no | no |
| CP37b | severe | F | 43 | Italian | 142 | | | | | | | | | | | | |
| CP38 | severe | F | 89 | Italian | 11 | fever, cough, dyspnea | death | | yes | yes | yes | no | no | no | no | no | no |
| CP39 | critical | M | 37 | Italian | 35 | fever, cough, dyspena | alive | | no | no | no | no | no | no | no | no | no |
| CP40 | severe | F | 67 | Italian | 19 | fever, cough | alive | | no | no | no | no | no | no | no | no | yes |
| CP41 | severe | M | 57 | Italian | 10 | fever, cough, diarrhea | alive | | no | no | yes | no | no | no | no | no | yes |
| CP42 | severe | M | 77 | Italian | 19 | fever, dyspnea | alive | | no | yes | yes | yes | no | no | no | no | no |
| CP43 | critical | M | 69 | Italian | 13 | fever, cough, dyspena | alive | | no | no | yes | yes | no | no | no | no | no |
| CP44a | severe | M | 60 | Italian | 16 | fever, dyspena | alive | | no | no | no | no | no | no | no | no | yes |
| CP44b | severe | M | 60 | Italian | 215 | | | | | | | | | | | | |
| CP45a | mild | M | 53 | Italian | 24 | fever | alive | | na | na | na | na | na | na | na | na | na |
| CP45b | mild | M | 53 | Italian | 204 | | | | | | | | | | | | |
| CP46b | severe | F | 80 | Italian | 67 | | | | | | | | | | | | |
| CP47a | severe | F | 59 | Italian | 26 | fever, dyspnea | alive | | no | no | no | no | no | no | yes | no | yes |
| CP48a | severe | F | 60 | Italian | 39 | fever cough, dyspnea | alive | | no | yes | yes | no | no | no | no | no | yes |
| CP48b | severe | F | 60 | Italian | 57 | | | | | | | | | | | | |
| CP49a | severe | M | 50 | Italian | 21 | fever, cough | alive | | no | no | no | no | no | no | no | no | no |
| CP49b | severe | M | 50 | Italian | 42 | | | | | | | | | | | | |
| CP50a | severe | M | 71 | Italian | 21 | fever, dyspnea | alive | | no | no | yes | no | no | no | no | no | yes |
| CP50b | severe | M | 71 | Italian | 42 | | | | | | | | | | | | |
| CP51 | critical | M | 55 | Italian | 26 | fever, dyspnea | alive | | no | no | no | no | no | no | no | no | no |
| CP52a | severe | M | 73 | Italian | 41 | fever, cough | alive | | no | no | no | no | no | yes | no | no | yes |
| CP52b | severe | M | 73 | Italian | 79 | | | | | | | | | | | | |
| CP53a | severe | M | 61 | Italian | 18 | fever, cough, dyspnea | alive | | no | no | yes | no | no | no | no | no | no |
| CP53b | severe | M | 61 | Italian | 72 | | | | | | | | | | | | |
| CP54a | moderate | M | 66 | Italian | 14 | fever, cough, dyspena | alive | | no | no | no | no | no | no | no | no | yes |
| CP54b | moderate | M | 66 | Italian | 77 | | | | | | | | | | | | |
| CP55a | severe | F | 76 | Italian | 14 | fever, cough | alive | | no | no | yes | no | no | yes | no | no | no |
| CP55b | severe | F | 76 | Italian | 84 | | | | | | | | | | | | |
| CP56a | severe | M | 69 | Italian | 14 | cough, dyspena, asthenia | alive | | no | no | no | no | yes | no | no | no | yes |
| CP56b | severe | M | 69 | Italian | 78 | | | | | | | | | | | | |
| CP57a | severe | F | 72 | Italian | 22 | fever, cough, dyspnea | alive | | no | no | yes | no | no | no | no | no | yes |
| CP57b | severe | F | 72 | Italian | 108 | | | | | | | | | | | | |
| CP58a | severe | M | 74 | Italian | 33 | fever, cough | alive | | yes | no | yes | no | no | no | no | no | yes |
| CP58b | severe | M | 74 | Italian | 109 | | | | | | | | | | | | |
| CP59 | mild | F | 37 | Italian | 18 | fever | alive | | na | na | na | na | na | na | na | na | na |
| CP60a | severe | F | 54 | Italian | 7 | fever, dyspnea, dyharrea, anosmia | alive | | no | no | no | no | no | no | no | no | no |
| CP60b | severe | F | 54 | Italian | 86 | | | | | | | | | | | | |
| CP61 | severe | M | 43 | Italian | 118 | fever, dyspnea | alive | | no | no | no | no | no | no | no | no | yes |
| CP62 | moderate | M | 49 | Italian | 110 | cough | alive | | no | no | no | yes | no | no | yes | no | yes |
| CP63 | severe | M | 32 | Italian | 85 | dyspnea | alive | | no | no | no | no | no | no | no | no | yes |
| CP64 | severe | F | 51 | Italian | 107 | fever, cough | alive | | no | no | no | no | no | no | no | no | no |
| CP65a | severe | M | 65 | Italian | 64 | fever | alive | | no | no | no | no | no | no | no | no | no |
| CP65b | severe | M | 65 | Italian | 176 | | | | | | | | | | | | |
| CP65C | severe | M | 65 | Italian | 288 | | | | | | | | | | | | |
| CP66 | severe | M | 56 | Italian | 93 | fever, dyspnea | alive | | no | no | no | no | yes | no | yes | no | yes |
| CP67 | severe | M | 52 | Italian | 89 | fever, dyspnea, anosmia | alive | | no | no | no | yes | no | no | yes | no | no |
| CP68a | moderate | M | 64 | Italian | 101 | fever, cough, dyspnea | alive | | no | no | no | no | no | no | no | no | no |
| CP68b | moderate | M | 64 | Italian | 210 | | | | | | | | | | | | |
| CP69 | severe | F | 56 | Italian | 112 | fever, cough, dyspnea | alive | | no | no | yes | no | no | no | no | no | no |
| CP70 | severe | M | 52 | Italian | 122 | fever, cough, hypoxia | alive | | no | no | yes | no | no | no | no | no | no |
| CP71 | mild | M | 29 | Swedish | 170 | myalgia, headache, anosmia, fatigue,chest pain | alive | | no | na | na | na | na | na | na | na | na |
| CP72a | mild | M | 65 | Swedish | 60 | fever | alive | | no | na | na | na | na | na | na | na | na |
| CP72b | mild | M | 65 | Swedish | 180 | | | | no | na | na | na | na | na | na | na | na |
| CP73 | mild | M | 48 | Swedish | 180 | fever, cough | alive | | no | na | na | na | na | na | na | na | na |
| CP73b | mild | M | 48 | Swedish | 330 | | | | no | na | na | na | na | na | na | na | na |
| CP74 | mild | F | 61 | Swedish | 198 | myalgia, cough, anosmia | alive | | no | na | na | na | na | na | na | na | na |
| CP74b | mild | F | 61 | Swedish | 393 | | | | no | na | na | na | na | na | na | na | na |
| CP75 | mild | M | 56 | Swedish | 193 | fatigue, myalgia, cough, headache, anosmia | alive | | no | na | na | na | na | na | na | na | na |
| CP75b | mild | M | 56 | Swedish | 388 | | | | no | na | na | na | na | na | na | na | na |
| CP76 | mild | F | 29 | Swedish | 219 | cough, anosmia | alive | | no | na | na | na | na | na | na | na | na |
| CP76b | mild | F | 30 | Swedish | 452 | | | | no | na | na | na | na | na | na | na | na |
| CP77 | mild | F | 43 | Swedish | 238 | fatigue, throat pain | alive | | no | na | na | na | na | na | na | na | na |
| CP78 | mild | M | 55 | Swedish | 157 | myalgia, fatigue, anosmia | alive | | no | na | na | na | na | na | na | na | na |
| CP79 | mild | F | 52 | Swedish | 160 | myalgia, fatigue, anosmia | alive | | no | na | na | na | na | na | na | na | na |
| CP81 | moderate | F | 88 | Italian | 11 | fever | alive | yes | no | no | yes | no | no | no | yes | no | yes |
| CP82 | severe | F | 79 | Italian | 15 | fever, dyspnea | alive | yes | no | no | yes | yes | no | yes | no | no | yes |
| CP83 | moderate | F | 48 | Italian | 7 | fever | alive | yes | no | no | yes | no | no | no | no | no | yes |
| CP84 | moderate | M | 83 | Italian | 15 | asthenia, diarrhea | alive | yes | no | yes | yes | yes | no | no | no | no | yes |
| CP85 | moderate | F | 53 | Italian | 10 | fever, asthenia | alive | no | no | no | no | no | no | no | no | no | no |
| CP86 | severe | F | 73 | Italian | 10 | fever | alive | no | no | no | no | no | no | no | no | no | no |
| CP87 | mild | M | 34 | Italian | 196 | fever, asthenia, anosmia | alive | no | no | no | no | no | no | no | no | no | no |
| CP87b | mild | M | 34 | Italian | 435 | | | | | | | | | | | | |
| CP88 | moderate | F | 81 | Italian | 26 | fever, asthenia | alive | yes | no | yes | no | yes | no | no | no | no | yes |
| CP89 | severe | M | 57 | Italian | 21 | fever, dyspnea | alive | yes | no | no | yes | yes | no | no | no | no | yes |
| CP90 | severe | F | 73 | Italian | 22 | fever, cough | alive | yes | no | yes | yes | yes | no | no | no | no | yes |
| CP91 | mild | M | 66 | Italian | 206 | fever, asthenia | alive | no | no | no | no | no | no | no | no | no | no |
| CP92 | severe | M | 80 | Italian | 9 | fever, asthenia, cough | alive | yes | no | no | yes | no | no | no | no | no | yes |
| CP93 | mild | F | 24 | Swedish | 188 | anosmia | alive | | no | na | na | na | na | na | na | na | na |
| CP94 | mild | F | 26 | Swedish | 191 | cough, anosmia | alive | | no | na | na | na | na | na | na | na | na |
| CP95 | mild | F | 24 | Swedish | 196 | fever | alive | | no | na | na | na | na | na | na | na | na |
| CP95b | mild | F | 25 | Swedish | 393 | | | | | | | | | | | | |
| CP96 | mild | F | 32 | Swedish | 218 | fever | alive | | no | na | na | na | na | na | na | na | na |
| CP97 | mild | F | 32 | Swedish | 229 | fever, cough | alive | | no | na | na | na | na | na | na | na | na |
| CP98 | mild | F | 59 | Swedish | 231 | cough | alive | | no | na | na | na | na | na | na | na | na |
| CP99 | mild | M | 30 | Swedish | 216 | fever, cough | alive | | no | na | na | na | na | na | na | na | na |
| CP99b | mild | M | 31 | Swedish | 381 | | | | | | | | | | | | |
| CP100 | mild | M | 39 | Swedish | 229 | fever, anosmia | alive | | no | na | na | na | na | na | na | na | na |
| CP101 | mild | F | 22 | Swedish | 69 | fever | alive | | no | na | na | na | na | na | na | na | na |
| CP102 | mild | M | 30 | Swedish | 205 | fever, anosmia | alive | | no | na | na | na | na | na | na | na | na |
| CP103 | mild | M | 22 | Swedish | 263 | anosmia | alive | | no | na | na | na | na | na | na | na | na |
| CP105 | mild | F | 33 | Swedish | 36 | fever, anosmia | alive | | no | na | na | na | na | na | na | na | na |
| CP105b | mild | F | 33 | Swedish | 140 | | | | | | | | | | | | |
| CP106 | mild | F | 30 | Swedish | 32 | fever, cough, anosmia | alive | | no | na | na | na | na | na | na | na | na |
| CP106b | mild | F | 30 | Swedish | 165 | | | | | | | | | | | | |

| New ID | other disease or comorbidities (if other comorb yes) | antiviral 1:ribavirin 2:lopinavir/ritonavir-hydroxicloroquine 3: hydroxicloroquine 4:remdesivir | date antiviral therapy | steroid | antibiotics | Oxigen therapy | ICU |
|---|---|---|---|---|---|---|---|
| CP01 | | 3/4 | 23-3-2020 | yes | yes (not known) | VM 40% | no |
| CP02 | | 2 | 2020-03-09 | no | Piperacillin/tazobactam, doxycyclin | VM 60% | no |
| CP03a | | 2 | 2020-03-08 | no | levofloxacin/ceftriaxone Piperacillin/tazobactam, doxycyclin | CPAP | no |
| CP03b | | | | | | | |
| CP03c | | | | | | | |
| CP03d | | | | | | | |
| CP04 | emophilia A | 1 | 2020-03-02 | no | piperacillin/tazobactam | HFNC | no |
| CP06a | | 2 | 2020-03-05 | no | Piperacillin/tazobactam/ levofloxacin | VM 50% | no |
| CP06b | | | | | | | |
| CP06c | | | | | | | |
| CP07a | | na | na | na | na | no | no |
| CP07b | | | | | | | |
| CP08a | | 2 | 2020-03-07 | no | Amoxicillin/clavulanic acid doxycyclin | no | no |
| CP08b | | | | | | | |
| CP08c | | | | | | | |
| CP09a | | 1/2 | 2020-02-25 | no | Piperacillin/tazobactam azithromycin vancomycin | no | no |
| CP09b | | | | | | | |
| CP09c | | | | | | | |
| CP10a | | no | / | no | no | no | no |
| CP10b | | | | | | | |
| CP11a | | no | / | no | no | no | no |
| CP11b | | | | | | | |
| CP12 | | 2 | 2020-03-02 | no | Piperacillin/tazobactam, doxycyclin | HFNC | no |
| CP13a | | 2 | 2020-03-06 | no | na | no | no |
| CP13b | | | | | | | |
| CP14 | previous colon carcinoma | 2 | 2020-02-27 | no | | VM 60% | no |
| CP15 | | 1 | 2020-03-13 | no | piperacillin/tazobactam | no | no |
| CP16a | | 3 | 2020-03-12 | no | Ceftriaxone/doxicyclin | VM 60% | no |
| CP16b | | | | | | | |
| CP17a | | 2 | 2020-03-07 | no | Piperacillin/tazobactam, doxycyclin | LFNC | no |
| CP17b | | | | | | | |
| CP18 | | 2 | 2020-03-02 | no | | VM 60% | no |
| CP19a | gatroesophageal reflux, renal disfunction | 1/2 | 2020-02-28 | no | Piperacillin/tazobactam, doxycyclin/ Vancomicina | HFNC | no |
| CP19b | | | | | | | |
| CP19c | | | | | | | |
| CP20 | | 2 | 2020-02-22 | no | Piperacillin/tazobactam, linezolid, vancomycin, azithromycin | VM 40% | no |
| CP21 | | 1/2 | 2020-02-28 | no | | VM 35% | no |
| CP23 | | 2 | 2020-03-21 | no | Ceftriaxone, azythromicin, doxycyclin, trimethoprim-sulfamethoxazole | HFNC | no |
| CP24 | | 3 | 2020-03-23 | no | Ceftriaxone/doxicyclin | HFNC | no |
| CP25 | osteoporosis, anemia, previous anticoagulant lupus | 4 | 2020-03-21 | no | trimethoprim-sulfamethoxazole | no | no |
| CP26 | | 2 | 2020-03-07 | no | | HFNC/CPAP | yes |
| CP27 | mammalian carcinoma | 2/4 | 2020-03-17 | no | | HFNC/CPAP | no |
| CP28 | microcytic anemia | 2/4 | nd | yes | Piperacillin/tazobactam, doxycyclin | HFNC/CPAP | no |
| CP29 | | 2 | 2020-03-24 | yes | ceftriaxone/doxicyclin azithromycin | HFNC/CPAP | yes |
| CP29b | | | | | | | |
| CP30 | hypothyroidism, previous prostatic carcinoma | 2 | 2020-03-25 | no | Piperacillin/tazobactam azithromycin | HFNC | no |
| CP31a | na | na | na | na | na | na | na |
| CP31b | | | | | | | |
| CP33 | | 2 | 2020-02-25 | no | Piperacillin/tazobactam, doxycyclin | CPAP | no |
| CP34a | previous pneumococcical pneumonia | 2 | 2020-02-26 | no | Piperacillin/tazobactam, doxycyclin | HFNC | no |
| CP34b | | | | | | | |
| CP34c | | | | | | | |
| CP35a | | 2 | 2020-03-08 | no | | HFNC | no |
| CP35b | | | | | | | |
| CP36 | | 2 | 2020-03-05 | no | piperacillin/tazobactam doxycyclin | no | no |
| CP37a | | 2 | 2020-02-28 | no | Piperacillin/tazobactam, doxycyclin | HFNC | no |
| CP37b | | | | | | | |
| CP38 | | 2 | 2020-03-11 | no | na | na | no |
| CP39 | | 2 | 2020-02-22 | yes | Piperacillin/tazobactam, trimethoprim-sulfamethoxazole | HFNC/CPAP | yes |
| CP40 | hypothyroidism | no | / | no | Piperacillin/tazobactam, doxycyclin | CPAP | no |
| CP41 | not specified | 3 | 2020-03-19 | no | ceftriaxone/azithromycin | CPAP | no |
| CP42 | | 2 | 2020-03-15 | yes | Piperacillin/tazobactam, doxycyclin, azithromycin | HFNC/CPAP | no |
| CP43 | | 3 | 2020-03-18 | no | Ceftriaxone/doxicyclin | CPAP | yes |
| CP44a | peripheral artherial disease | 3 | na | yes | ceftriaxone/azithromycin | HFNC | no |
| CP44b | | | | | | | |
| CP45a | | na | / | na | na | no | no |
| CP45b | | | | | | | |
| CP46b | | | | | | | |
| CP47a | depressive syndrome | 2 | 2020-03-13 | no | | HFNC | no |
| CP48a | not specified | 2 | 2020-03-05 | yes | Piperacillin/tazobactam | CPAP/PEEP | no |
| CP48b | | | | | | | |
| CP49a | | 3 | 2020-03-20 | yes | Piperacillin/tazobactam-azithromycin trimetropim | VM 40% | no |
| CP49b | | | | | | | |
| CP50a | prostatic hyperplasia | 3 | 2020-03-25 | yes | ceftriaxone/azithromycin | HFNC | no |
| CP50b | | | | | | | |
| CP51 | | 2 | 2020-03-17 | no | piperacillin/tazobactam, trimethoprim-sulfamethoxazole, doxicyclin | HFNC, CPAP | yes |
| CP52a | prostatic hyperplasia, gladbladder stones | 3 | 2020-04-06 | no | azithromycin | HFNC | no |
| CP52b | | | | | | | |
| CP53a | | 2 | 2020-03-07 | no | | CPAP | no |
| CP53b | | | | | | | |
| CP54a | not specified | no | / | no | ceftriaxone/azithromycin | no | no |
| CP54b | | | | | | | |
| CP55a | | 1/2 | 2020-02-28 | no | Ceftazidime/doxicyclin | LFNC | no |
| CP55b | | | | | | | |
| CP56a | psoriatic arthritis | 2 | na | no | Amoxicillin/clavulanic acid | LFNC | no |
| CP56b | | | | | | | |
| CP57a | dyslipidemia | 1/2 | 2020-02-27 | no | | VM 60% | no |
| CP57b | | | | | | | |
| CP58a | Paget syndrome | 2 | 2020-02-27 | no | Piperacillin/tazobactam, doxycyclin | CPAP | no |
| CP58b | | | | | | | |
| CP59 | | na | na | na | na | no | no |
| CP60a | | 3 | 2020-03-29 | yes | Ceftriaxone/amoxicillin azithromycin | VM | no |
| CP60b | | | | | | | |
| CP61 | gastroesophageal reflux, kidney stones | 3 | 2020-03-27 | yes | yes (not known) | VM/HFNC, CPCP | no |
| CP62 | ovarian cystis and uterine fibroid | no | / | yes | | no | no |
| CP63 | gastritis (pump inhinitors treatment) | 3 | 2020-04-03 | no | broad spectrum antibiotics | HFNC | no |
| CP64 | | 3 | na | no | yes (not known) | HFNC | no |
| CP65a | | 3 | 2020-04-23 | no | Ceftriaxone/doxicyclin | VM/ CPAP | no |
| CP65b | | | | | | | |
| CP65C | | | | | | | |
| CP66 | hypothyroidism | 3 | 2020-03-30 | yes | ampicillin/sulbactam | PEEP/CPAP- VM | no |
| CP67 | | 3 | na | yes | ceftriaxone/azithromycin | HFNC | no |
| CP68a | | 3 | 2020-03-22 | yes | Ceftriaxone/doxicyciclin Piperacillin/tazobactam azithromycin | no | no |
| CP68b | | | | | | | |
| CP69 | | 2 | 2020-03-11 | no | Trimethoprim-sulfamethoxazole, doxicyclin, piperacillin/tazobactam, azithromycin | CPAP | no |
| CP70 | | 3 | 2020-04-01 | yes | ceftriaxone/azithromycin | VM | no |
| CP71 | na | na | na | na | na | na | na |
| CP72a | na | na | na | na | na | na | na |
| CP72b | na | na | na | na | na | na | na |
| CP73 | na | na | na | na | na | na | na |
| CP73b | na | na | na | na | na | na | na |
| CP74 | na | na | na | na | na | na | na |
| CP74b | na | na | na | na | na | na | na |
| CP75 | na | na | na | na | na | na | na |
| CP75b | na | na | na | na | na | na | na |
| CP76 | na | na | na | na | na | na | na |
| CP76b | na | na | na | na | na | na | na |
| CP77 | na | na | na | na | na | na | na |
| CP78 | na | na | na | na | na | na | na |
| CP79 | na | na | na | na | na | na | na |
| CP81 | dyslipidemia, renal cystis, osteoporosis, stroke | no | / | | no | no | no |
| CP82 | ostheoarthritis | 4 | 2020-09-08 | no | doxicyclin | VM60% | no |
| CP83 | hypothiroidism, diverticulitis (sigmoid) | no | / | no | amoxicillin/clavulanic acid | no | no |
| CP84 | chronic anemia, cronic renal disease | no | / | no | vancomycin | no | no |
| CP85 | | no | / | no | doxicyclin | no | no |
| CP86 | | no | / | yes | ceftriaxone/azythromicin | HFNC | no |
| CP87 | | 2 | 2020-03-23 | no | azythromicin | no | no |
| CP87b | | | | | | | |
| CP88 | transient ischemic attack | no | / | no | no | no | no |
| CP89 | renal disfunction | no | / | yes | Ceftriaxone | VM40% | no |
| CP90 | Sjogren syndrom, previous B lymphoma | no | / | yes | Ceftriaxone | HFNC | no |
| CP91 | | 2 | 2020-03-15 | no | no | no | no |
| CP92 | renal disease, depression | no | / | no | yes (not known) | VM | no |
| CP93 | na | na | na | na | na | na | na |
| CP94 | na | na | na | na | na | na | na |
| CP95 | na | na | na | na | na | na | na |
| CP95b | | | | | | | |
| CP96 | na | na | na | na | na | na | na |
| CP97 | na | na | na | na | na | na | na |
| CP98 | na | na | na | na | na | na | na |
| CP99 | na | na | na | na | na | na | na |
| CP99b | | | | | | | |
| CP100 | na | na | na | na | na | na | na |
| CP101 | na | na | na | na | na | na | na |
| CP102 | na | na | na | na | na | na | na |
| CP103 | na | na | na | na | na | na | na |
| CP105 | na | na | na | na | na | na | na |
| CP105b | | | | | | | |
| CP106 | na | na | na | na | na | na | na |
| CP106b | | | | | | | |

| New ID | chest CT scan | pneumonia severity | complications | laboratory data collection | C-reactive protein (CRP) (<0,5) mg/L) | pO2 | pO2/FIO2 ratio | haemoglobin (g/dL) | platelets (cells/ul) | WBC (cells/ul) | neutro (cells/ul) | lymph (cells/ul) | urea (mg/dl) | creatine (mg/dl) | ALT (mU/mL) |
|---|---|---|---|---|---|---|---|---|---|---|---|---|---|---|---|
| CP01 | interstitial pneumonia (low, bilateral) | | renal complication | 2020-03-14 | 2,73 | 58,2 | 277 | 17,8 | 173000 | 7300 | 6100 | 600 | 89 | 1,02 | 19 |
| CP02 | interstitial tickness lobe of the right lung | | no | 2020-02-29 | 1,25 | 70,1 | 334 | 12,7 | 178000 | 4200 | 3000 | 600 | 54 | 0,73 | 24 |
| CP03a | interstitial lung disease (low, bilateral) | | no | 2020-03-05 | 3,68 | na | na | 14,7 | 244000 | 6100 | 4300 | 1100 | 26 | 0,92 | 47 |
| CP03b | | | | | | | | | | | | | | | |
| CP03c | | | | | | | | | | | | | | | |
| CP03d | | | | | | | | | | | | | | | |
| CP04 | na | | na | 2020-03-02 | 5,97 | na | na | 15 | 132000 | 5600 | 3500 | 1500 | 33 | 0,89 | 33 |
| CP06a | bronchovascular thickening and opacity | | no | 2020-03-04 | 20,39 | na | na | 14,7 | 245000 | 7900 | 5800 | 900 | 42 | 0,79 | 15 |
| CP06b | | | | | | | | | | | | | | | |
| CP06c | | | | | | | | | | | | | | | |
| CP07a | na | | no | na | na | na | na | na | na | na | na | na | na | na | na |
| CP07b | | | | | | | | | | | | | | | |
| CP08a | lung opacity basal bilateral | | no | 2020-03-06 | 3,38 | na | na | 12,8 | 97000 | 3500 | 1900 | 1200 | 21 | 0,52 | 20 |
| CP08b | | | | | | | | | | | | | | | |
| CP08c | | | | | | | | | | | | | | | |
| CP09a | bronchovascular thickening | | no | 2020-02-25 | 7,5 | na | na | 13,8 | 290000 | 5800 | 4100 | 1200 | 29 | 0,85 | 13 |
| CP09b | | | | | | | | | | | | | | | |
| CP09c | | | | | | | | | | | | | | | |
| CP10a | no lung densification | | no | 2020-02-27 | 0,77 | na | na | 12 | 163000 | 4000 | 2900 | 700 | 26 | 0,63 | 29 |
| CP10b | | | | | | | | | | | | | | | |
| CP11a | no lung densification | | no | 2020-02-22 | 0,55 | na | na | 12,9 | 242000 | 4100 | 2400 | 100 | 27 | 0,76 | 12 |
| CP11b | | | | | | | | | | | | | | | |
| CP12 | bronchovascular thickening | | no | 2020-03-01 | 6,49 | na | na | 15,3 | 284000 | 10900 | 8380 | 1250 | 33 | 1,11 | 38 |
| CP13a | na | | na | 2020-03-06 | 14,75 | 52,9 | 106 | 11,6 | 441000 | 9400 | 7200 | 1300 | 40 | 0,73 | 53 |
| CP13b | | | | | | | | | | | | | | | |
| CP14 | interstitial lung disease | | no | 2020-02-28 | 5,05 | na | na | 14,2 | 138000 | 3500 | 2500 | 800 | 44 | 0,84 | 22 |
| CP15 | na | | na | 2020-03-12 | 15,62 | na | na | 13,4 | 269000 | 13560 | 12140 | 730 | 200 | 1,51 | 31 |
| CP16a | lung densification | | no | 2020-03-12 | 13,1 | 120 | 120 | 13,9 | 275000 | 6600 | 4700 | 1300 | 29 | 0,59 | 35 |
| CP16b | | | | | | | | | | | | | | | |
| CP17a | bronchovascular thickening | | no | 2020-03-07 | 2,8 | na | na | 16,3 | 203000 | 4500 | 3000 | 900 | 33 | 0,9 | 41 |
| CP17b | | | | | | | | | | | | | | | |
| CP18 | bilateral densification (medium-low area) | | no | 2020-02-29 | 16,68 | 53,1 | 253 | 14,8 | 229000 | 8600 | 7500 | 600 | 42 | 1,01 | 57 |
| CP19a | bilateral densification | | no | 2020-02-28 | 6,13 | 59,3 | 283 | 15,6 | 180000 | 5200 | 4100 | 700 | 51 | 1,5 | 33 |
| CP19b | | | | | | | | | | | | | | | |
| CP19c | | | | | | | | | | | | | | | |
| CP20 | lung densification (low, bilateral) | | no | 2020-02-22 | 5,79 | na | na | 14,4 | 141000 | 3600 | 2600 | 800 | 37 | 0,77 | 39 |
| CP21 | multiple bilateral densification | | no | 2020-02-25 | 2,84 | na | na | 15,1 | 148000 | 4500 | 2500 | 1600 | 49 | 1,2 | 30 |
| CP23 | bilateral lung densification | | no | 2020-03-21 | 15,14 | 65,5 | 312 | 14,5 | 110000 | 5100 | 3700 | 1000 | na | 0,77 | 86 |
| CP24 | multiple bilateral densification | | no | 2020-03-19 | 21,84 | 128 | 128 | 15 | 206000 | 9080 | 8080 | 440 | 33 | 0,72 | 59 |
| CP25 | interstitial lung densification (peripheral basal area, right) | | no | 2020-03-21 | 2,10 | na | na | 11 | 174000 | 5100 | 4000 | 600 | 29 | 1,23 | 48 |
| CP26 | bronchovascular thickening | | dx foot swelling | 2020-03-07 | 12,56 | 74,6 | 213 | 12,7 | 164000 | 4800 | 3800 | 700 | 24 | 0,71 | 138 |
| CP27 | bilateral interstitial opacity | | no | 2020-03-13 | 14,14 | 73,2 | 73,2 | 12,8 | 223000 | 5700 | 4600 | 600 | 42 | 0,74 | 33 |
| CP28 | irregular densification | | no | 2020-03-13 | 4,82 | 58 | 276 | 8 | 325000 | 5200 | 2370 | 1100 | na | 0,65 | 19 |
| CP29 | bronchovascular thickening | | no | 2020-03-23 | 26,35 | 76,5 | 128 | 13,2 | 287000 | 13400 | 11700 | 600 | 39 | 0,73 | 55 |
| CP29b | | | | | | | | | | | | | | | |
| CP30 | bilateral densification (basal area) | | anemia | 2020-03-25 | 14,55 | 55,7 | 265 | 11,1 | 239000 | 7100 | 5600 | 800 | 36 | 0,82 | 54 |
| CP31a | na | na | na | na | na | na | na | na | na | na | na | na | na | na | na |
| CP31b | | | | | | | | | | | | | | | |
| CP33 | lung opacity | | no | 2020-02-26 | 12,24 | na | na | 15,3 | 214000 | 5500 | 3200 | 1300 | 37 | 1,19 | 28 |
| CP34a | parenchimal densification (low basal area left) | | no | 2020-02-25 | 2,93 | na | na | 12,3 | 103000 | 2800 | 1800 | 600 | 26 | 0,68 | 38 |
| CP34b | | | | | | | | | | | | | | | |
| CP34c | | | | | | | | | | | | | | | |
| CP35a | bilateral interstitial opacity | | no | 2020-03-05 | 20,45 | 72 | 343 | 14,3 | 318000 | 6610 | 5400 | 550 | 20 | 0,78 | 60 |
| CP35b | | | | | | | | | | | | | | | |
| CP36 | na | | na | 2020-03-04 | 11,50 | na | na | 12,4 | 254000 | 8260 | 7320 | 400 | 42 | 0,64 | 46 |
| CP37a | bronchovascular thickening | | no | 2020-02-27 | 11,16 | na | na | 12,5 | 299000 | 4500 | 4000 | 400 | 21 | 0,66 | 58 |
| CP37b | | | | | | | | | | | | | | | |
| CP38 | na | | na | 2020-03-16 | na | 72 | 72 | 11,9 | 246000 | 18700 | 10540 | 13800 | 108 | 1,2 | 26 |
| CP39 | bronchovascular thickening | | sugar rising | 2020-02-22 | 19,70 | na | na | 13,2 | 143000 | 5500 | 4700 | 300 | 59 | 1,14 | 21 |
| CP40 | interstitial lung disease and parenchimal densification (left), | | no | 2020-03-09 | na | 89.3 | 94,8 | 12,6 | 314000 | 12300 | 10900 | 400 | 48 | 0,54 | 15 |
| CP41 | na | | na | 2020-03-23 | 14,40 | na | na | 13,5 | 148000 | 5000 | 3580 | 1210 | 19 | 0,81 | 60 |
| CP42 | interstitial lung disease (medium-low, bilateral) | | skin rash | 2020-03-14 | 14,98 | 83,1 | 805 | 15,9 | 131000 | 4000 | 2900 | 800 | 104 | 1,61 | 82 |
| CP43 | interstitial lung disease | | no | 2020-03-18 | 11,68 | 77,3 | 77,3 | 16,6 | 172000 | 6000 | 4500 | 900 | 37 | 0,69 | 35 |
| CP44a | bilateral lung opacity | | no | 2020-03-21 | 17,64 | 57,7 | 275 | 15,9 | 179000 | 8400 | 6900 | 900 | 40 | 0,74 | 146 |
| CP44b | | | | | | | | | | | | | | | |
| CP45a | na | | no | na | na | na | na | na | na | na | na | na | na | na | na |
| CP45b | | | | | | | | | | | | | | | |
| CP46b | | | | | | | | | | | | | | | |
| CP47a | lung opacity | | no | 2020-03-13 | 11,35 | 62,4 | 297 | 12,3 | 189000 | 4400 | 3300 | 900 | 23 | 0,72 | 128 |
| CP48a | lung opacity (medium left area) | | no | 2020-03-05 | 38,96 | 47,2 | 225 | 16,7 | 215000 | 14400 | 13100 | 800 | 60 | 1,17 | 21 |
| CP48b | | | | | | | | | | | | | | | |
| CP49a | bilateral densification | | no | 2020-03-17 | 14,42 | 96,8 | 242 | 13,5 | 180000 | 6400 | 5300 | 600 | 35 | 0,82 | 18 |
| CP49b | | | | | | | | | | | | | | | |
| CP50a | lung bilateral densification | | no | 2020-03-24 | 11,93 | 58,7 | 280 | 12,9 | 299000 | 6800 | 5600 | 700 | 37 | 0,77 | 40 |
| CP50b | | | | | | | | | | | | | | | |
| CP51 | multiple lung densification | | *C.parapsilosis* | 2020-03-02 | 14,70 | na | na | 13,3 | 137000 | 3900 | 2900 | 800 | 34 | 0,97 | 130 |
| CP52a | parenchimal densification (right lung, low area) | | no | 2020-04-05 | 2,41 | 73,2 | 349 | 13,8 | 187000 | 6400 | 4900 | 100 | 31 | 0,72 | 21 |
| CP52b | | | | | | | | | | | | | | | |
| CP53a | lung densification | | no | 2020-03-10 | 3,69 | 150 | 250 | 14 | 313000 | 7000 | 5900 | 800 | 37 | 0,86 | 57 |
| CP53b | | | | | | | | | | | | | | | |
| CP54a | na | | na | 2020-03-10 | 3,49 | 98,8 | 250 | 15,5 | 201000 | 7000 | 5200 | 800 | na | na | na |
| CP54b | | | | | | | | | | | | | | | |
| CP55a | lung opacity (medium-low left region) | | no | 2020-02-28 | 1,15 | na | na | 13,8 | 205000 | 3540 | 2620 | 560 | 46 | 1,56 | 29 |
| CP55b | | | | | | | | | | | | | | | |
| CP56a | bronchovascular thickening (medium-low right) | | no | 2020-03-07 | 17,16 | 60,9 | 290 | 14,2 | 265000 | 9500 | 8000 | 100 | 65 | 0,83 | 17 |
| CP56b | | | | | | | | | | | | | | | |
| CP57a | multiple lung foci | | no | 2020-02-27 | 13,43 | na | na | 11,6 | 185000 | 3100 | 2400 | 500 | 18 | 0,61 | 37 |
| CP57b | | | | | | | | | | | | | | | |
| CP58a | densification medium-high left lung, opacitity medium-high right lung | | no | 2020-02-27 | 9,37 | na | na | 14,7 | 153000 | 4100 | 2800 | 800 | 39 | 0,85 | 42 |
| CP58b | | | | | | | | | | | | | | | |
| CP59 | na | | no | na | na | na | na | na | na | na | na | na | na | na | na |
| CP60a | bilateral parenchimal opacity | | no | 2020-03-29 | 6,1 | 64,8 | 309 | 13,3 | 103000 | 2100 | 1200 | 700 | 15 | 0,55 | 119 |
| CP60b | | | | | | | | | | | | | | | |
| CP61 | lung densification (basal right and left) | | skin rash | 2020-02-29 | 3,32 | 64,6 | 308 | 15,5 | 118000 | 2300 | 1400 | 600 | 42 | 0,97 | 19 |
| CP62 | lung opacity (bilateral) | | no | 2020-03-11 | 21,36 | 69,2 | 330 | 12,6 | 205000 | 5300 | 3700 | 1200 | na | 0,39 | 42 |
| CP63 | bronchovascular thickening | | no | 2020-04-03 | 12,72 | 63,9 | 305 | 14,5 | 257000 | 9140 | 7820 | 710 | 19 | 0,82 | 23 |
| CP64 | lung densification | | no | 2020-03-16 | 11,77 | 75,4 | 359 | 11,9 | 244000 | 4860 | 4060 | 500 | na | 0,52 | na |
| CP65a | bilateral lung opacity and initial lung densification | | na | 2020-04-23 | 12,43 | 58 | 276 | 15,2 | 219000 | 6020 | 4090 | 1160 | 37 | 1,19 | 13 |
| CP65b | | | | | | | | | | | | | | | |
| CP65C | | | | | | | | | | | | | | | |
| CP66 | densification medium-low right and low left lung | | B.fragilis, C.parapsilosis (wound swab) | 2020-03-30 | 13,35 | 81,7 | 81,7 | 9,5 | 264000 | 5900 | 3800 | 900 | 62 | 3,23 | 20 |
| CP67 | pleural effusion (basal right lung) | | no | 2020-04-02 | 3,49 | 66,2 | 315 | 14,3 | 142000 | 5510 | 2520 | 2380 | 37 | 0,64 | 40 |
| CP68a | na | | na | 2020-03-20 | 14,07 | 61,6 | 294 | 14,7 | 236000 | 8800 | 6900 | 1300 | 32 | 0,7 | 48 |
| CP68b | | | | | | | | | | | | | | | |
| CP69 | bronchovascular thickening | | cerebellar ischemia | 2020-03-11 | 32,86 | 119 | 198 | 13,9 | 244000 | 9500 | 8100 | 900 | 77 | 1,31 | 14 |
| CP70 | medium-upper right and medium left parenchiamal densification | | no | 2020-03-31 | 8,96 | 29,2 | 314 | 14,3 | 155000 | 4700 | 3700 | 700 | 34 | 0,95 | 39 |
| CP71 | na | na | na | na | na | na | na | na | na | na | na | na | na | na | na |
| CP72a | na | na | na | na | na | na | na | na | na | na | na | na | na | na | na |
| CP72b | na | na | na | na | na | na | na | na | na | na | na | na | na | na | na |
| CP73 | na | na | na | na | na | na | na | na | na | na | na | na | na | na | na |
| CP73b | na | na | na | na | na | na | na | na | na | na | na | na | na | na | na |
| CP74 | na | na | na | na | na | na | na | na | na | na | na | na | na | na | na |
| CP74b | na | na | na | na | na | na | na | na | na | na | na | na | na | na | na |
| CP75 | na | na | na | na | na | na | na | na | na | na | na | na | na | na | na |
| CP75b | na | na | na | na | na | na | na | na | na | na | na | na | na | na | na |
| CP76 | na | na | na | na | na | na | na | na | na | na | na | na | na | na | na |
| CP76b | na | na | na | na | na | na | na | na | na | na | na | na | na | na | na |
| CP77 | na | na | na | na | na | na | na | na | na | na | na | na | na | na | na |
| CP78 | na | na | na | na | na | na | na | na | na | na | na | na | na | na | na |
| CP79 | na | na | na | na | na | na | na | na | na | na | na | na | na | na | na |
| CP81 | no | | no | 2020-09-10 | 1,79 | na | na | 12,9 | 161000 | 5800 | 3723 | 1235 | na | 0,95 | 10 |
| CP82 | bilateral densification low-medium area | | no | 2020-09-05 | 2,86 | na | na | 12,3 | 178000 | 3640 | 1950 | 1360 | 38 | 0,98 | 18 |
| CP83 | absence of evident lesions | | no | 2020-09-15 | 6,43 | na | na | 13,3 | 253000 | 8960 | 7220 | 1060 | na | 0,77 | 28 |
| CP84 | bronchovascular thickening, dx basal lung opacity | | *C difficile* | 2020-09-15 | 3,83 | na | na | 10,2 | 218000 | 5380 | 4200 | 620 | 76 | 1,78 | 21 |
| CP85 | multiple bilateral densification | | no | 2020-09-21 | 2,06 | na | na | 14,6 | 230000 | 5450 | 3000 | 1800 | 21 | 0,75 | 15 |
| CP86 | initial lung densification | | no | 2020-09-20 | 4,51 | na | na | 12,2 | 151000 | 5050 | 3830 | 900 | 25 | 0,76 | 14 |
| CP87 | no | | no | na | na | na | na | na | na | na | na | na | na | na | na |
| CP87b | | | | | | | | | | | | | | | |
| CP88 | no lung densification | | no | 2020-09-26 | 0,19 | 75,1 | 358 | 12,6 | 420000 | 9200 | 7600 | 1000 | 17 | 0,45 | 22 |
| CP89 | multiple bilateral densification | | no | 2020-09-20 | 12,25 | na | na | 14,9 | 170000 | 10800 | 8380 | 1300 | 97 | 1,22 | 38 |
| CP90 | bilateral densification (basal area) | | | 2020-09-10 | 2,66 | 87,4 | 124 | 12,2 | 186000 | 8400 | 7400 | 500 | 47 | 1,09 | 22 |
| CP91 | no | | no | na | na | na | na | na | na | na | na | na | na | na | na |
| CP92 | multiple bilateral densification | | no | 2020-10-10 | 15,38 | na | na | 12 | 206000 | 8490 | 6980 | 670 | 45 | 1,2 | 17 |
| CP93 | na | na | na | na | na | na | na | na | na | na | na | na | na | na | na |
| CP94 | na | na | na | na | na | na | na | na | na | na | na | na | na | na | na |
| CP95 | na | na | na | na | na | na | na | na | na | na | na | na | na | na | na |
| CP95b | | | | | | | | | | | | | | | |
| CP96 | na | na | na | na | na | na | na | na | na | na | na | na | na | na | na |
| CP97 | na | na | na | na | na | na | na | na | na | na | na | na | na | na | na |
| CP98 | na | na | na | na | na | na | na | na | na | na | na | na | na | na | na |
| CP99 | na | na | na | na | na | na | na | na | na | na | na | na | na | na | na |
| CP99b | | | | | | | | | | | | | | | |
| CP100 | na | na | na | na | na | na | na | na | na | na | na | na | na | na | na |
| CP101 | na | na | na | na | na | na | na | na | na | na | na | na | na | na | na |
| CP102 | na | na | na | na | na | na | na | na | na | na | na | na | na | na | na |
| CP103 | na | na | na | na | na | na | na | na | na | na | na | na | na | na | na |
| CP105 | na | na | na | na | na | na | na | na | na | na | na | na | na | na | na |
| CP105b | | | | | | | | | | | | | | | |
| CP106 | na | na | na | na | na | na | na | na | na | na | na | na | na | na | na |
| CP106b | | | | | | | | | | | | | | | |

| | | | | | | | | | | | | | | | | | | | | | | | | | | | | | | | | | | | | | | | |
|---|---|---|---|---|---|---|---|---|---|---|---|---|---|---|---|---|---|---|---|---|---|---|---|---|---|---|---|---|---|---|---|---|---|---|---|---|---|---|---|
| **CP107** | mild | M | 69 | Swedish | 63 | fever | alive | | no | na | na | na | na | na | na | na | na | na | na | na | na | na | na | na | na | na | na | na | na | na | na | na | na | na | na | na | na | na | na |
| **CP108** | mild | M | 67 | Swedish | 63 | fever, anosmia | alive | | no | na | na | na | na | na | na | na | na | na | na | na | na | na | na | na | na | na | na | na | na | na | na | na | na | na | na | na | na | na | na |
| **CP123b** | mild | M | 66 | Italian | 106 | fever | alive | no | no | no | no | no | no | no | no | no | no | | no | no | no | no | no | no | no | | no | 2020-11-12 | na | na | na | na | na | na | na | na | na | na | na |
| **CP162** | mild | M | 45 | Swedish | 86 | fever, asthenia | alive | | no | na | na | na | na | na | na | na | na | na | na | na | na | na | na | na | na | na | na | na | na | na | na | na | na | na | na | na | na | na | na |
| **CP163** | mild | F | 43 | Swedish | 266 | fever, cough | alive | | no | na | na | na | na | na | na | na | na | na | na | na | na | na | na | na | na | na | na | na | na | na | na | na | na | na | na | na | na | na | na |
| **CP164** | mild | F | 54 | Swedish | 74 | fever, cough, anosmia, myalgi | alive | | no | na | na | na | na | na | na | na | na | na | na | na | na | na | na | na | na | na | na | na | na | na | na | na | na | na | na | na | na | na | na |
| **CP164b** | mild | F | 54 | Swedish | 171 | | | | | | | | | | | | | | | | | | | | | | | | | | | | | | | | | | |
| **CP166** | mild | M | 56 | Swedish | 70 | fever | alive | | no | na | na | na | na | na | na | na | na | na | na | na | na | na | na | na | na | na | na | na | na | na | na | na | na | na | na | na | na | na | na |
| **CP167** | mild | F | 61 | Swedish | 271 | fever, cough | alive | | no | na | na | na | na | na | na | na | na | na | na | na | na | na | na | na | na | na | na | na | na | na | na | na | na | na | na | na | na | na | na |
| **CP167b** | mild | F | 61 | Swedish | 368 | | | | | | | | | | | | | | | | | | | | | | | | | | | | | | | | | | |
| **CP168** | mild | F | 41 | Swedish | 259 | na | alive | | no | na | na | na | na | na | na | na | na | na | na | na | na | na | na | na | na | na | na | na | na | na | na | na | na | na | na | na | na | na | na |
| **CP169** | mild | F | 50 | Swedish | 54 | cough | alive | | no | na | na | na | na | na | na | na | na | na | na | na | na | na | na | na | na | na | na | na | na | na | na | na | na | na | na | na | na | na | na |
| **CP170** | mild | F | 52 | Swedish | 31 | fever, cough | alive | | no | na | na | na | na | na | na | na | na | na | na | na | na | na | na | na | na | na | na | na | na | na | na | na | na | na | na | na | na | na | na |
| **CP170b** | mild | F | 52 | Swedish | 128 | | | | | | | | | | | | | | | | | | | | | | | | | | | | | | | | | | |
| **CP171** | mild | M | 56 | Swedish | 150 | fever, anosmia | alive | | no | na | na | na | na | na | na | na | na | na | na | na | na | na | na | na | na | na | na | na | na | na | na | na | na | na | na | na | na | na | na |
| **CP172** | mild | F | 55 | Swedish | 159 | na | alive | | no | na | na | na | na | na | na | na | na | na | na | na | na | na | na | na | na | na | na | na | na | na | na | na | na | na | na | na | na | na | na |
| **CP173** | mild | M | 66 | Swedish | 103 | fever | alive | | no | na | na | na | na | na | na | na | na | na | na | na | na | na | na | na | na | na | na | na | na | na | na | na | na | na | na | na | na | na | na |
| **CP175** | mild | M | 37 | Swedish | 440 | na | alive | | no | na | na | na | na | na | na | na | na | na | na | na | na | na | na | na | na | na | na | na | na | na | na | na | na | na | na | na | na | na | na |
| **CP176** | mild | F | 18 | Swedish | 446 | fever, anosmia | alive | | no | na | na | na | na | na | na | na | na | na | na | na | na | na | na | na | na | na | na | na | na | na | na | na | na | na | na | na | na | na | na |
| **CP177** | severe | M | 80 | Italian | 20 | fever,cough,pneomunia | alive | no | yes | no | yes | no | no | no | no | no | no | | no | / | yes | ceftriaxone, azythromicin, doxycyclin | VM | no | bronchovascular thickening | | no | 2021-03-09 | 4,07 | na | na | 11,5 | 219000 | 10800 | 9900 | 400 | 82 | 0,76 | 57 |
| **CP178** | severe | M | 77 | Italian | 27 | fever,cough,dyspnea | alive | no | no | no | no | no | no | no | no | no | no | | no | / | yes(dexamethasone, methylprisolone) | azythromicin | CPAP60%, HFNC, VM | no | ground glass lesions (medium lateral area), | | no | 2021-03-08 | 2,68 | 40,9 | 195 | 11,4 | 152000 | 8360 | 6590 | 1100 | 74 | 1,07 | 118 |
| **CP179** | severe | F | 67 | Italian | 15 | fever | alive | yes | no | no | no | yes | no | no | no | no | no | | no | / | yes | | VM, LFNC | no | no | | no | 2021-03-10 | 1,05 | 64,8 | 308 | 14,8 | 224000 | 5100 | 3600 | 600 | 29 | 0,53 | 18 |
| **CP180** | severe | M | 80 | Italian | 24 | | | | | | | | | | | | | | | | | | | | | | | | | | | | | | | | | | |
| **CP181** | severe | M | 86 | Italian | 15 | asthenia | alive | yes | yes | no | yes | no | no | no | no | no | no | | no | / | yes (dexamethasone) | ceftriaxone/doxicyclin | CPAP, HFNC | no | interstitial thickening | | no | 2021-03-11 | 16,99 | 74,4 | 186 | 13,5 | 240000 | 12000 | 10100 | 900 | 78 | 1,2 | 47 |
| **CP182** | severe | F | 75 | Italian | 24 | asthenia | alive | no | no | no | no | no | no | no | no | no | no | | no | / | yes (dexamethasone) | no | CPAP, HFNC, VM | no | ground glass lesions (low area) bilateral densification | | no | 2021-03-10 | 5,74 | na | na | 12,6 | 121000 | 3200 | 2800 | 300 | 116 | na | 34 |
| **CP183** | severe | M | 83 | Italian | 24 | fever, cough | alive | no | no | no | yes | no | no | no | no | no | no | dysplipidemia | no | / | | cefriaxone | CPAP60% | no | no | | no | 2021-03-07 | 22,57 | na | na | 13,8 | 175000 | 12600 | 11800 | 500 | 109 | 1,65 | 35 |
| **CP184** | severe | M | 72 | Italian | 23 | fever, cough, dyspnea | alive | yes | no | no | no | yes | no | yes | no | no | no | | hyperimmune plasma | 2021-03-15 | yes (dexamethasone) | ceftriaxone/doxicyclin | CPAP60% | no | bilateral densification (basal area) | | no | 2021-03-09 | 7,32 | 62,4 | 297 | 13,3 | 144000 | 7920 | 6420 | 780 | 30 | 0,84 | 26 |
| **CP185** | severe | M | 77 | Italian | 15 | fever, | alve | yes | yes | no | yes | no | no | no | no | no | no | | hyperimmune plasma | 2021-03-12 | yes (dexamethasone) | ceftriaxone | CPAP60%,VM | no | ground glass lesions (upper area) | | no | 2021-03-11 | 4,25 | 112 | 112 | 11,7 | 108000 | 3040 | 2700 | 240 | 130 | 4,58 | 28 |
| **CP186** | severe | F | 70 | Italian | 24 | fever, cough | alive | yes | no | no | yes | no | no | no | no | no | no | | no | / | yes (dexamethasone) | ceftriaxone | CPAP60%,VM60% | no | bilateral densification | | no | 2021-03-12 | 9,62 | na | na | 12,5 | 122000 | 4300 | 3500 | 600 | 56 | 1,1 | 32 |
| **CP187** | severe | M | 51 | Italian | 40 | fever, dyspnea | alive | yes | no | no | yes | no | no | no | no | no | no | | no | / | yes (dexamethasone) | ceftriaxone | CPAP60%,VM,HFNC | no | | | no | 2021-03-03 | 4,28 | 51 | 243 | 13,5 | 259000 | 8200 | 7400 | 200 | 52 | 1,17 | 63 |
| **CP188** | severe | F | 83 | Italian | 34 | fever, cough, dyspnea | alive | yes | no | no | yes | no | no | no | no | no | no | | hyperimmune plasma | 2021-03-03 | yes(dexamethasone, methylprisolone) | Piperacillin/tazobactam | CPAP60%,VM60% | no | ground glass lesions | | no | 2021-03-01 | 0,08 | 55,5 | 264 | 12,9 | 139000 | 7360 | 5360 | 1380 | 68 | 1,09 | 32 |
| **CP189** | mild | M | 81 | Italian | 22 | fever, dyspnea | alive | yes | no | no | yes | no | yes | no | no | no | no | | nd | nd | yes | yes | nd | nd | nd | | nd | 2021-03-10 | 2,14 | na | na | 12,2 | 146000 | 5800 | 4300 | 1100 | 68 | 1,22 | 24 |
| **CP190** | severe | M | 78 | Italian | 21 | fever, cough, dyspnea | alive | yes | no | no | yes | no | no | no | no | no | yes | kidney failure | no | / | yes (dexamethasone) | ceftriaxone | CPAP,PEEP | no | ground glass lesions | | no | 2021-03-12 | 9,14 | 64,2 | 306 | 15,1 | 202000 | 8150 | 6720 | 880 | 67 | 1,51 | 31 |
| **CP191** | severe | F | 61 | Italian | 27 | fever | alive | yes | no | no | yes | yes | yes | no | no | no | yes | CMV pneumonia infection (08/2019),TEP, BPCO,lung graft | 1, hyperimmune plama | 05/03/2021(ribavirine) 12/03/2021(hyperimmune plasma) | yes (dexamethasone) | | VM (40%) | no | ground glass lesions | | no | 2021-03-04 | 3,93 | na | na | 10,2 | 73000 | 4600 | 3400 | 1000 | 79 | 1,91 | 24 |
| **CP192** | severe | F | 57 | Italian | 26 | fever | alive | yes | yes | no | yes | no | yes | no | no | no | yes | lung graft | hyperimmune plasma | 2021-03-02 | yes (dexamethasone, prednisone)) | Piperacillin/tazobactam | LFNC(2L/min) | no | ground glass lesions | | no | 2021-03-01 | 2,2 | na | na | 10,5 | 176000 | 4200 | 2700 | 800 | 162 | 2,63 | 16 |
| **CP193** | mild | M | 31 | Italian | 32 | fever (post vaccine infection) | alive | no | no | no | no | no | no | no | no | no | no | | no | | no | | no | | | | | na | na | na | na | na | na | na | na | na | na | na | na |
| **CP194** | moderate | F | 22 | Italian | 17 | cough, asthenia, dyspnea | alive | yes | no | no | no | no | no | no | no | no | no | GPA PR3+ pANCA, previous SARS-CoV-2 infection in July 2020 | no | / | prednisone | trimethoprim/sulfamethoxazole | no | no | intestitial addensation | | | 2021-03-27 | 4,34 | na | na | 9,7 | 614000 | 8660 | 6740 | 1170 | 29 | 0,71 | 24 |
| **CP196** | critical | M | 62 | Italian | 22 | fever, dysgueusia, cough | alive | na | na | na | na | na | na | na | na | na | na | | hyperimmune plasma | 2021-02-16 | no | no | CPAP/VM60% | yes | intestitial bilateral pneumonia | | copies/mL | 2021-02-15 | 12,43 | na | na | 14,5 | 165000 | 6200 | 4600 | 1100 | 59 | 1,04 | 26 |

**M** **Male**
**F** **Female**
**na** **Not available**
**hcv** **Hepatitis C virus**
**VM** **Venturi oxygen mask**
**CPAP** **Continuous positive airway pressure**
**HFNC** **High-flow nasal cannula**
**ICU** **Intensive care unit**
**CT** **Computed tomography**
